# A codon-sensitive conformational switch gates commitment to translation start sites

**DOI:** 10.64898/2026.03.24.713952

**Authors:** Sydney F. McGuire, Matthew C. Chan, Tung Ching Chan, Rachel R. Cueny, Niseema Pachikara, Eva M. Alleman, Veronica M. Sikora, Arvind Rasi Subramaniam, Melody G. Campbell, Christopher P. Lapointe

**Affiliations:** Basic Sciences Division, Fred Hutchinson Cancer Center, Seattle, WA; Computational Biology Section of Public Health Sciences Division, Fred Hutchinson Cancer Center, Seattle, WA

## Abstract

Human translation initiation requires single-nucleotide precision to establish the reading frame, yet initiation at non-AUG codons plays key roles in gene expression. How the initiation machinery balances precision with this regulated flexibility remains unclear. Here, we define a conformational branchpoint governed by the human initiation factor eIF5 that gates commitment to start codons. Using single-molecule, structural, and molecular dynamics approaches, we demonstrate that eIF5 reversibly occupies at least two conformations, which depends on a strictly conserved loop in the protein that monitors start codon identity. AUG codons favor the conformation that is stabilized by an eIF5-stimulated GTP hydrolysis step, which commits the complex to the start site. Non-AUG codons favor a ‘standby’ conformation that destabilizes eIF5 and may place its N-terminal domain at a site shared with eIF2β, its ancient structural homolog. This branchpoint complements upstream start codon selection steps and intrinsically controls the efficiency of non-AUG initiation.

---

To establish the reading frame for protein synthesis, the human translation initiation machinery must recognize the start codon (AUG) with single-nucleotide precision. However, non-AUG start codons (e.g., CUG) can also mediate protein synthesis, albeit with reduced efficiency^1–5^. While some messenger RNAs (mRNAs) rely solely on a non-AUG start codon to synthesize the encoded protein, many non-AUG sites have regulatory roles. Upstream open reading frames, non-canonical open reading frames, and 5’ extensions or truncations of proteins often use non-AUG start codons^5–8^. Differential usage of AUG versus non-AUG codons plays key roles during mitosis, meiosis, and cellular stress responses^5,8–10^. Dysregulated usage contributes to cancer progression and metastasis^11^ as well as neurological disorders^12–15^. The molecular events that underlie recognition of the start codon remain unclear.

Translation initiation is directed by eukaryotic initiation factors (eIFs) and the initiator methionyl-tRNA_i_^Met^ (tRNA_i_). The process starts when a 43S preinitiation complex – the small (40S) ribosomal subunit bound by eIF1, eIF1A, eIF3, and the eIF2–GTP–tRNA_i_ complex – loads onto the mRNA, which is facilitated by eIF4 proteins bound near the 5’-m^7^G cap^16–18^. While directionally moving 5’ to 3’ along the 5’UTR, the complex scans for translation start codons^19–21^, with selection enhanced by favorable Kozak sequence context^22^. A start codon correctly placed at the ribosomal P site and base paired with the anticodon of tRNA_i_ remodels the complex from a scanning-permissive conformation to a scanning-arrested one^19,20,23–25^. This transition triggers initial departure of eIF1. While paused at the putative start codon, eIF1 and eIF5 then directly compete for binding to drive opposite outcomes^26^. Stable eIF1 rebinding is favored at non-AUG codons, which may allow scanning to resume^27^. Successful eIF5 binding allows eIF5B to bind and direct joining of the large (60S) subunit to form the ribosome^28–32^.

eIF5 binding represents a key commitment step of initiation. The protein stimulates GTP hydrolysis by eIF2 (or inorganic P_i_ release)^33–36^ and commits the complex to the start site by initiating the cascade of events that completes formation of the ribosome^26^. Intriguingly, high levels (or activity) of eIF5 promote non-AUG initiation, whereas lower ones favor AUG codons^26,37–39^. These genetic effects may result entirely from the direct competition between eIF5 and eIF1 to bind initiation complexes once scanning arrests at candidate start codons^10,26^. Alternatively, eIF5 itself also may govern a downstream, codon-sensitive branchpoint that directly controls the efficiency of initiation.

To examine eIF5 directly, we combined single-molecule spectroscopy and cryo-EM using a reconstituted human translation initiation system, which we complemented with biochemical assays and molecular dynamics simulations. Our data reveal a conformational branchpoint governed by eIF5 upstream of GTP hydrolysis by eIF2 that requires a strictly conserved loop in eIF5 to monitor codon-anticodon base pairing. Collectively, our findings provide a biophysical framework that enhances understanding of how the initiation machinery balances single-nucleotide precision with regulated use of non-AUG start codons.

## RESULTS

### Direct analysis of eIF5 binding at start codons

Since eIF5 functions once initiation complexes pause at candidate start codons^26^, we reasoned that fidelity branchpoints governed by eIF5 may be masked in assays that include scanning and eIF1-dependent steps. To bypass those upstream steps, we leveraged the internal ribosome entry site encoded by hepatitis C virus (here-after, IRES) (**Extended Data Fig. 1A**). The IRES directly binds to the 40S ribosomal subunit and assembles the initiation complex precisely at the encoded AUG start codon, requiring only eIF2 (comprised of α, β, and γ subunits), tRNA_i_, and eIF1A ^40,41^. This IRES supports translation when programmed with non-AUG start codons in human cells^40^. Using single-particle cryo-EM, we confirmed that eIF5 occupied its canonical binding site^42–44^ on the IRES-assembled initiation complex (2.6 Å resolution); the N-terminal domain of eIF5 was positioned proximal to the eIF2-GDPNP-tRNA_i_ ternary complex and eIF1A, over-lapping the eIF1 binding site (**Fig. 1A,B, Extended Data Fig. 1B-H**, and **Supplementary Table 1**). Further consistent with prior structures^19,20,44,45^, the lack of density for the core domain of eIF2β indicated that this domain is released after recognition of the start codon but prior to GTP hydrolysis (since GDPNP was present) (**Fig. 1B**), likely with eIF2β maintained flexibly on the complex by contacts with eIF2γ^20,46,47^. Thus, the IRES represents a useful tool to isolate eIF5 function on initiation complexes paused at start codons.

**Figure 1:**
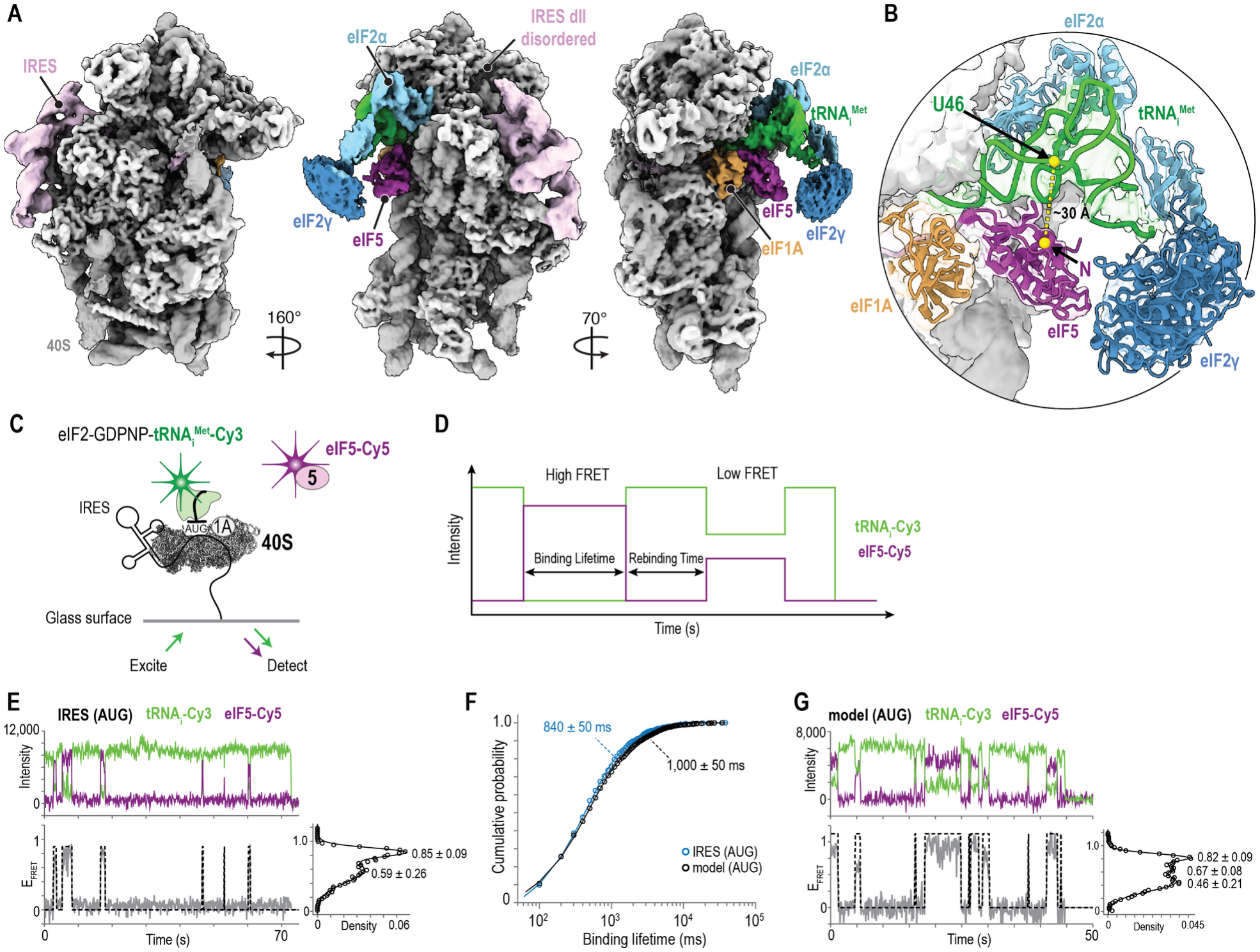
Direct analysis of eIF5 binding at start codons. **A**, Cryo-EM map of the initiation complex bound to the HCV IRES with an AUG start site. Density for eIF2γ is shown at a lower threshold. **B**, Close-up view of the N-terminal domain of eIF5 (purple) bound proximal to the tRNA_i_ (green). The labeling positions of U46 on tRNA_i_ (green) and the N terminus of eIF5 (purple) are indicated. **C**, Schematic of the single-molecule initiation assay. **D**, Theoretical single-molecule fluorescence data defining FRET states, binding lifetimes, and rebinding times. **E**, Example single-molecule fluorescence data where tRNA_i_–Cy3 (green) and eIF5–Cy5 (purple) were monitored. **F**, Cumulative probability plot of the eIF5 binding lifetime on the IRES and the model mRNA. Lines represent fits to a double-exponential function and the derived mean lifetimes are reported on the plot. **G**, Example single-molecule fluorescence data where tRNA_i_–Cy3 (green) and eIF5–Cy5 (purple) were monitored during canonical translation initiation on the model mRNA. In panels E and G, the dashed lines demarcate eIF5 binding events. The corresponding FRET efficiency distributions indicate the FRET means ± standard deviation. Distributions include only frames during eIF5 binding events.

To analyze eIF5 directly, we designed a single-molecule Förster resonance energy transfer (FRET) assay. As the FRET donor, we labeled synthetic tRNA_i_ on U46 with a Cy3 fluorescent dye (tRNA_i_-Cy3). We also labeled recombinant eIF5 on its N-terminus using a ybbR tag and Cy5 dye (FRET acceptor)^26^. This labeling position had no measurable impact on eIF5 function in vitro^26^ and should place the dyes within about 30 Å in initiation complexes (**Fig. 1B**). We assembled complexes that contained 40S subunits, 3’-biotinylated IRES, eIF1A, and the eIF2-GDPNP-tRNA_i_-Cy3 ternary complex, and we tethered them to a neutravidin-coated glass surface (**Fig. 1C**). We imaged in the presence of eIF5-Cy5 using total internal reflection fluorescence microscopy at 100 ms temporal resolution (**Extended Data Fig. 2A** and **Supplementary Table 2**).

eIF5 bound initiation complexes formed on the IRES similar to canonical complexes. As predicted, we observed eIF5 binding and dissociation as pulses of tRNA_i_-Cy3 (green) to eIF5-Cy5 (purple) FRET (**Fig. 1D,E**). eIF5 remained bound for 840 ± 50 ms on average (**Fig. 1F**). The protein bound repeatedly to individual complexes, as expected given our use of a non-hydrolyzable GTP analog (GDPNP). The time between eIF5 binding events decreased as the concentration of eIF5 increased (30 ± 1 µM^-1^s^-1^) (**Extended Data Fig. 2B, C**), which confirmed that eIF5 fully dissociated between binding events^26^. Importantly, we observed almost identical eIF5 binding lifetimes on canonical complexes formed on a model unstructured mRNA in the presence of eIF1 and eIF3 (**Fig. 1F, G**, and **Extended Data Fig. 2D,E**).

Together, our cryo-EM and single-molecule findings indicate that the IRES allows direct examination of eIF5 at start codons, independent of scanning and competition with eIF1. The broad FRET distributions observed here on both RNAs (**Fig. 1E,G**) and in a prior study^26^ indicate that eIF5-bound initiation complexes adopt multiple conformations. Whether these conformations represent static subpopulations or dynamically interconvert on a rapid timescale (<100 ms) remained unclear.

### Rapid and reversible sampling of multiple conformations

To resolve rapid conformational dynamics, we examined eIF5 binding at 5-fold higher temporal resolution using single-molecule FRET. The enhanced resolution was achieved by replacing the Cy5 dye on eIF5 with LD655 dye (FRET acceptor)^48,49^, which improved the signal and anti-correlation between donor and acceptor dyes when collecting data using 20 ms exposures (**Extended Data Fig. 3A-D**). During individual binding events using the LD655 dye, we observed rapid and strongly anti-correlated changes in tRNA_i_ and eIF5 fluorescence intensities (median Pearson r < -0.8) (**Fig. 2A** and **Extended Data Fig. 3E-F**). These FRET fluctuations represent dynamic repositioning of tRNA_i_ and eIF5 relative to one another within the complex throughout the binding event (**Fig. 2B**).

**Figure 2:**
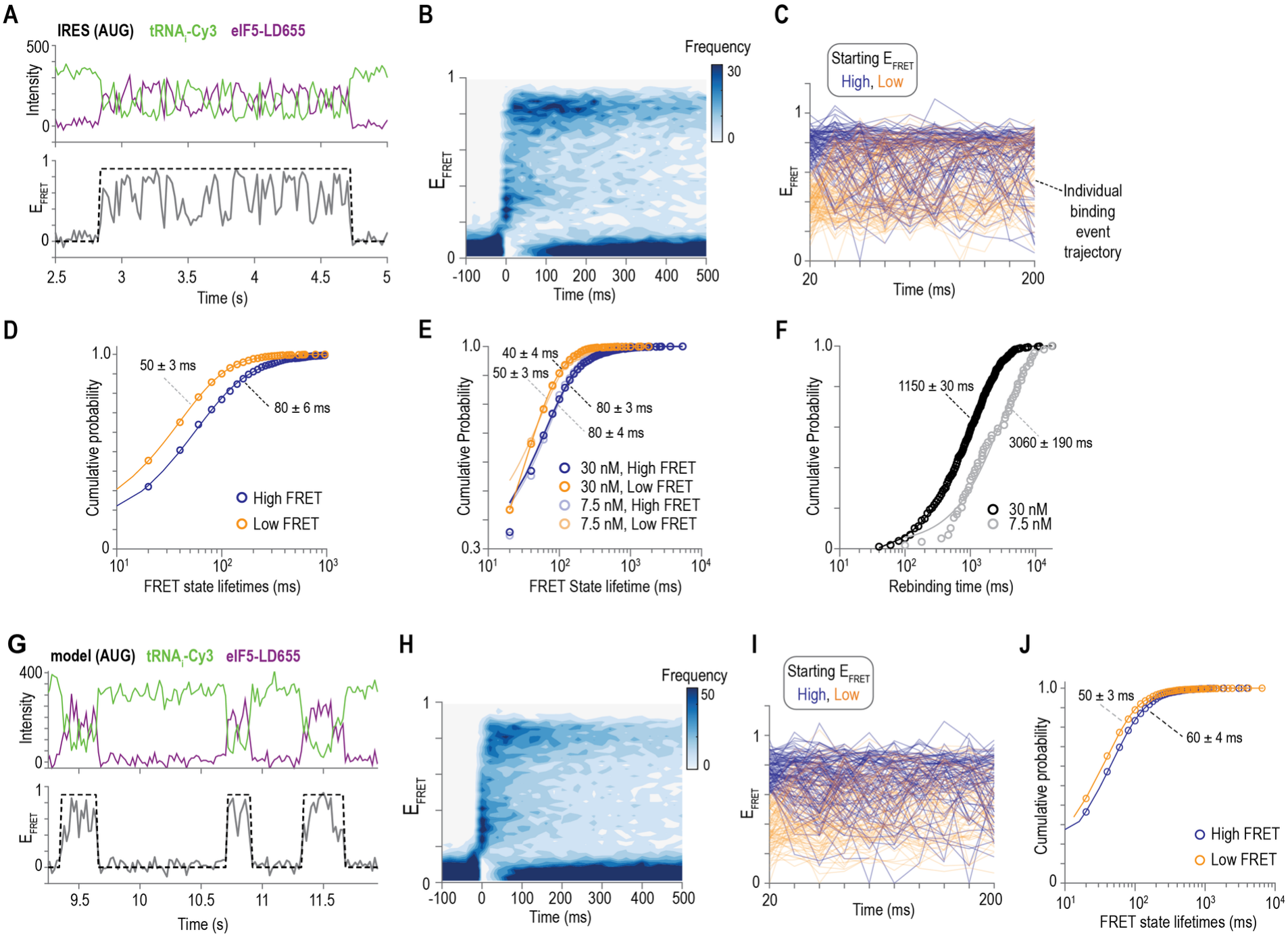
Rapid and reversible sampling of multiple conformations. **A**, Example single-molecule fluorescence data collected at 20 ms temporal resolution with tRNAi–Cy3 (green) and eIF5–LD655 (purple) on the IRES complex. **B**, Density plot of the observed FRET Efficiency (E_FRET_) synchronized to the start of all eIF5 binding events (716) observed on 314 complexes. **C**, Plot of the observed FRET efficiency trajectories of two hundred binding events over the first two hundred milliseconds. Blue lines represent events that started in high FRET (≥ 0.66) and orange lines represent events that started in low FRET (≤ 0.65). **D-E**, Cumulative probability plots of the high- and low-FRET state lifetimes with 5 nM eIF5 present (panel D), and 7.5 nM or 30 nM present (panel E). **F**, Cumulative probability plot of the eIF5 rebinding time at the indicated concentrations. **G-J**, Depict the same plots as in panels A-D but for eIF5 binding on canonical initiation complexes with an AUG start site in the presence of eIF1 and eIF3. Panel H depicts 1,318 binding events across 387 complexes. In panels A and G, the dashed lines demarcate the eIF5 binding events. In panels D, E, F, and J, the lines represent fits to exponential functions and the derived mean times ± 95 % CI are reported.

We analyzed the dynamics during individual events. To retain single-molecule insights, we overlaid the FRET efficiencies (E_FRET_) from the first 200 ms of individual binding events, which were color coded according to the minimum between the low-FRET and high-FRET populations (E_FRET_ = 0.65) (**Fig. 2C** and **Extended Data Fig. 3G**). As highlighted by the rapid intermixing of the trajectories, 86 % (476 / 553) of eIF5 binding events contained at least one transition between high- and low-FRET states. Using these threshold-based definitions, we estimated a high-FRET state lifetime of 80 ± 6 ms and a low-FRET state lifetime of 50 ± 3 ms during individual binding events (**Fig. 2D**). Adjustments to the threshold yielded highly similar lifetime distributions (**Extended Data Fig. 3H**). Furthermore, the high- and low-FRET state lifetimes were identical at different concentrations of eIF5 (**Fig. 2E,F**), indicative of intramolecular dynamics. The 14% of binding events (77 / 553) without a fluctuation were approximately 2-fold briefer than events with one and may represent complexes where eIF5 departed prior to a fluctuation or potential off-pathway complexes. Regardless, our findings demonstrate that eIF5-bound initiation complexes fluctuate between distinct conformations on a tens-of-milliseconds timescale during individual binding events.

The rapid tRNA_i_-to-eIF5 FRET fluctuations could represent generalizable dynamics of initiation complexes or arise from IRES-specific features. To distinguish these possibilities, we re-examined eIF5 binding to canonical initiation complexes formed on the model unstructured mRNA in the presence of eIF1 and eIF3. Importantly, the eIF5-bound complex exhibited rapid and reversible FRET fluctuations during individual binding events (**Figs. 2G-J** and **Extended Data Fig. 3I-L**). The means and lifetimes of the two conformations were nearly identical to those on the IRES. As additional validation, we examined eIF5 binding on a version of the IRES that lacked domain II, which can modulate the conformation of the 40S ribosomal subunit but disengages once eIF2-GTP-tRNA_i_ binds (**Fig. 1A** and **Extended Data Fig. 1A**) ^30,50–52^. While deletion of domain II from the IRES reduced eIF5 binding by 7-fold, the relatively rare eIF5 binding events that did occur maintained rapid fluctuations between high- and low-FRET states with lifetimes nearly identical to the wild-type IRES and model mRNA (**Extended Data Fig. 4**).

Together, our findings indicate that eIF5-bound initiation complexes rapidly and reversibly sample at least two conformations when paused at start codons, independent of IRES-specific features. Since the conformations interconvert during individual eIF5 binding events, they may correspond to distinct functional states of the initiation complex during start codon recognition.

### Start codon identity biases conformational occupancy

We asked whether the conformational dynamics of eIF5-bound initiation complexes depend on the identity of the start codon. In our cryo-EM structure, an asparagine residue of eIF5 (N_30_) resides within 3 Å of the interaction between U35 in the tRNA_i_ anticodon and the first nucleotide of the start codon (**Fig. 3A**). This residue and its flanking glycines (G_29_ and G_31_) form a loop in eIF5 strictly conserved from protists to humans (**Fig. 3B**). Identical contacts between eIF5 and the codon-anticodon interface were observed in published structures of canonical initiation complexes^42–44^. Furthermore, a foundational yeast genetic study linked this same loop to start codon fidelity^53^.

**Figure 3:**
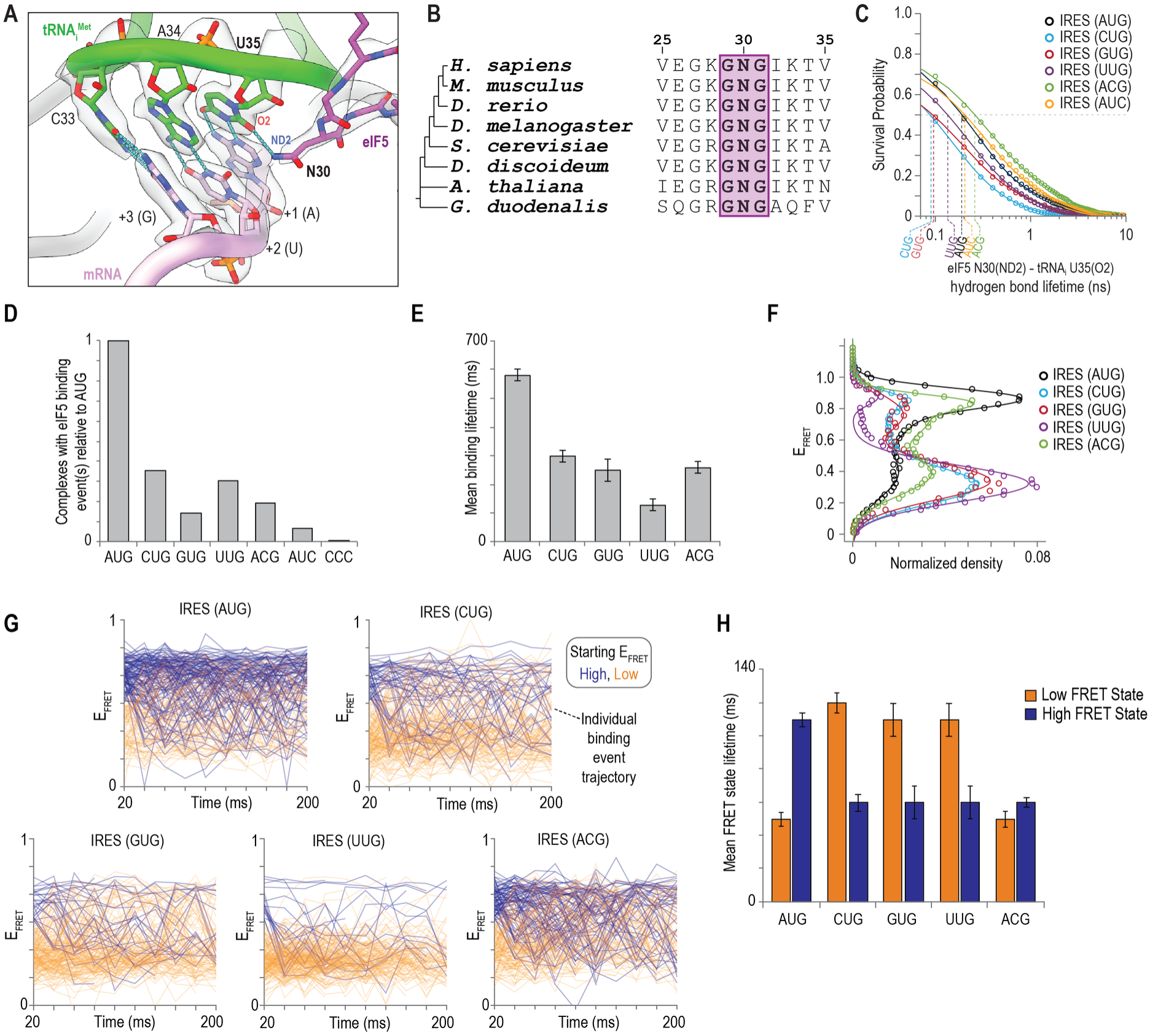
Start codon identity biases conformational occupancy. **A**, Structure of the eIF5 G_29_N_30_G_31_ loop bound ∼3 Å from the first nucleotide of the translation start site. The grey surface represents the electron density map, with local resolution of 2.8 Å. **B**, Multiple sequence alignment highlighting the strictly conserved G_29_N_30_G_31_ loop of eIF5. **C**, Survival probability plots of eIF5 N_30_ -tRNA_i_ U35 hydrogen bond lifetime at the indicated start site variants derived from 2.3 µs all atom molecular dynamics simulations. Dotted lines represent the computed halftime of the N_30_-U35 hydrogen bond. **D**, Fraction of initiation complexes bound by eIF5 at different start sites relative to binding at the canonical AUG start codon. **E**, Weighted population means of eIF5 binding lifetimes at different translation start sites. **F**, tRNA_i_-Cy3 – to – eIF5-LD655 FRET efficiency distributions observed at the indicated IRES start site variants; the lines represent fits to double gaussian functions. **G**, Plots of the observed FRET efficiency trajectories of two hundred binding events over the first two hundred milliseconds. Blue lines represent events that started in high FRET (≥ 0.66) and orange lines represent events that started in low FRET (≤ 0.65). **H**, Weighted population means of the high- and low-FRET state lifetimes at the indicated translation start sites.

Consistent with a role at the codon-anticodon interface, molecular dynamics simulations revealed that the amide side chain of N_30_ in eIF5 forms hydrogen bonds with the pyrimidine oxygen atom of U35 in the anticodon of tRNA_i_ (**Fig. 3C** and **Extended Data Fig. 5**). Optimal A-U base pairing facilitates stable N_30_ interactions, with a codon-anticodon RMSD of <1 Å and a hydrogen bond donor-acceptor distance of 3.5 Å (**Extended Data Fig. 5D**). In contrast, non-AUG codons disrupt the base pairing geometry, destabilizing N_30_ hydrogen bonding interactions, as demonstrated by shifts in the conformational landscape of the codon-anticodon interface (**Extended Data Fig. 5E-F**). Specifically, substitution of the first nucleotide in the start codon reduced the lifetime of the N_30_-U35 hydrogen bond interaction to different extents. Substitutions at the second or third nucleotide had minimal impact, but fewer hydrogen-bonding events occurred (**Fig. 3C, Supplementary Table 3**). Thus, all non-AUG codons we simulated rewired the N_30_ hydrogen bonding network (**Extended Data Fig. 5G**).

We hypothesized that alteration of local interactions at the codon-anticodon interface reshape the conformational ensemble of eIF5-bound initiation complexes. To test this model, we examined eIF5 on initiation complexes programmed with non-AUG codons using our single-molecule FRET assay (**Extended Data Fig. 6A-D**). CUG, UUG, GUG, and ACG start sites reduced the frequency and duration of eIF5 binding events to different extents (**Fig. 3D, E** and **Extended Data Fig. 6E**). Relative to AUG, the CUG codon had the least pronounced impact on eIF5 binding events (3-fold reduction) and their duration (300 ± 20 ms, 2-fold briefer), consistent with CUG displaying the highest translation efficiency in cells^54^. Substitution of the third position (AUC) nearly eliminated eIF5 binding (15-fold reduction), comparable to the IRES variant that lacked a recognizable start site (CCC) (**Fig. 3D**). In parallel with these kinetic effects, substitution of the first nucleotide shifted the FRET efficiency distributions almost entirely to the low-FRET conformation, due to 2-fold stabilization of the low-FRET state and 2-fold destabilization of the high-FRET state (**Fig. 3F-H** and **Extended Data Fig. 6F**). Substitution of the second nucleotide selectively destabilized the high-FRET state. Thus, non-AUG start codons both destabilized eIF5 binding and increased relative occupancy of the low-FRET conformation.

Our findings demonstrate that single-nucleotide mis-matches at the interface between the start codon and the anticodon of tRNA_i_ reshape the local interactions of N_30_ and ultimately the conformational dynamics of eIF5-bound initiation complexes. AUG codons favor the high-FRET conformation, whereas non-AUG codons favor the low-FRET one. While less efficiently bound, presumably due to leaky scanning, we observed similar effects using canonical initiation complexes assembled on an unstructured model mRNA with a CUG start site (**Extended Data Fig. 7**), corroborating our findings on the IRES. How the presence of a codon-anticodon mismatch biased residence in the two distinct conformations remained unclear.

### A conserved eIF5 loop controls codon sensitivity

We hypothesized that the single-nucleotide sensitivity of eIF5 dynamics required its G_29_N_30_G_31_ loop, given its position adjacent to the codon-anticodon interface and strict conservation (**Fig. 3A**). Consistent with this model, substitution of N_30_ with alanine reduced the duration of eIF5 binding events by 2-fold on initiation complexes poised at an AUG codon (**Fig. 4A**). This N30A variant also predominantly shifted residence to the low-FRET conformation, due to both stabilization of the low-FRET state (2fold) and destabilization of the high-FRET state (4-fold) (**Fig. 4B,C**). Similar effects occurred on complexes with the CUG start site (**Fig. 4A-C** and **Extended Data Fig. 8A**). Outside the loop, an R15M substitution in eIF5, which prevents GTP hydrolysis by eIF2γ^36^, retained rapid interconversions between the low- and high-FRET conformations, albeit with modest reductions in their lifetimes (**Fig. 4A-C** and **Extended Data Fig. 8A-C**). A K143E variant, which inverts a distal electrostatic contact with the 18S rRNA, also allowed rapid interconversion between the two conformations, despite a 3-fold reduction in the duration of the binding events (**Fig. 4A** and **Extended Data Fig. 8A,D**). Thus, substitution of N_30_ selectively biases eIF5-bound initiation complexes toward the low-FRET conformation.

**Figure 4:**
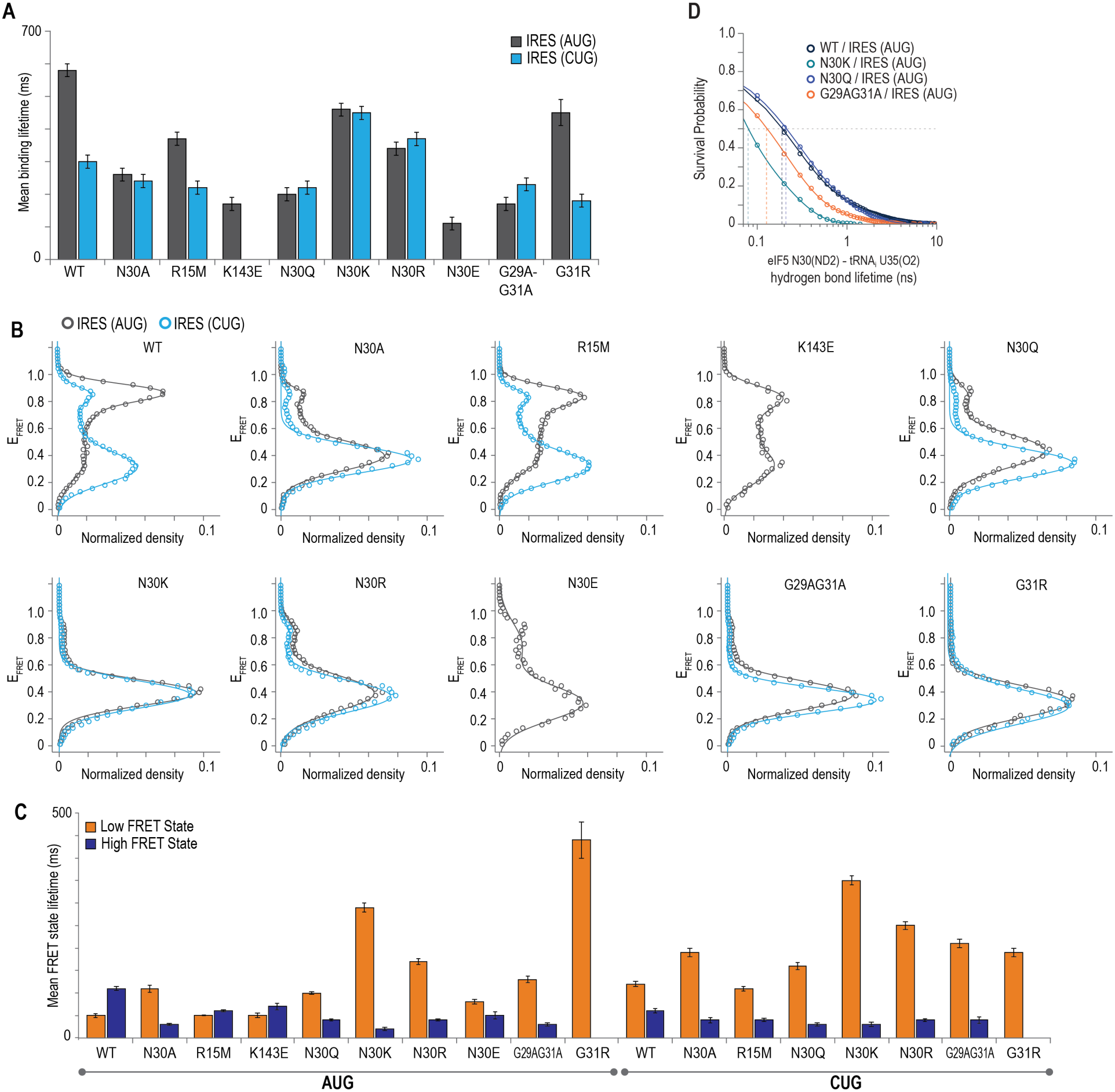
A conserved eIF5 loop controls codon sensitivity. **A**, Weighted population means of binding lifetimes for eIF5 variants at AUG and CUG start codons. **B**, tRNA_i_ -Cy3 – to – eIF5-LD655 FRET efficiency distributions observed for each eIF5 variant at AUG and CUG start sites; the lines represent fits to double gaussian functions. **C**, Weighted population means of the high- and low-FRET state lifetimes of each eIF5 variant at the AUG and CUG start sites. **D**, Survival probability plots of eIF5 N_30_-tRNA_i_ U35 hydrogen bond lifetimes for the indicated eIF5 variants derived from 2.3 µs all atom molecular dynamics simulations.

To understand the biochemical basis of the codon sensitivity, we examined how amino acid substitutions in the G_29_N_30_G_31_ loop affected eIF5 dynamics. Variants that altered side chain length, charge, and local backbone flexibility all shifted predominant occupancy to the low-FRET conformation (**Fig. 4B,C** and **Extended Data Fig. 8A**). Simply lengthening the side chain (N30Q) inhibited eIF5 binding, destabilized the high-FRET state, and stabilized the low-FRET state, each by 2–3 fold. Introduction of a positive charge (N30K, N30R) further enhanced occupancy of the low-FRET state, and a negative charge shortened binding events by 5-fold, consistent with electrostatic repulsion. Restriction of local backbone flexibility by substitution of the flanking glycines with alanine also strongly biased occupancy of the low-FRET state. A G31R variant essentially eliminated the high-FRET state. We observed similar effects on canonical initiation complexes that loaded and scanned for start codons in real time on an m^7^G capped mRNA in the presence of eIF4 (**Extended Data Fig. 9**). Wild-type eIF5 rapidly and reversibly sampled both low- and high-FRET states, and the G31R substitution nearly eliminated occupancy of the high-FRET state (**Extended Data Fig. 9**). Complementary molecular dynamics simulations moreover revealed reduced hydrogen-bonding potential between eIF5 loop variants and U35 in tRNA_i_ despite stable codon-anticodon geometry (**Fig. 4D** and **Extended Data Fig. 5**).

Together, these findings establish that interactions at the codon-anticodon interface transmit through the G_29_N_30_G_31_ loop to shape the conformational ensemble occupied by eIF5-bound initiation complexes. Multiple intrinsic properties of the loop allow occupancy of the high-FRET conformation on AUG codons. Substitutions within the G_29_N_30_G_31_ loop that altered side chain chemistry, length, charge, or backbone flexibility all dramatically shifted relative residence to the low-FRET conformation. The attenuated effects on CUG codons further support a model in which the G_29_N_30_G_31_ loop determines codon-dependent shifts in the conformational ensemble. However, whether this conformational bias underlies commitment to a start codon by the initiation complex remained unclear.

### GTP hydrolysis stabilizes the high-FRET conformation

During initiation, eIF5 stimulates GTP hydrolysis by eIF2γ to progress initiation complexes to ribosomal subunit joining. In all experiments above, we used a non-hydrolyzable GTP analog (GDPNP) to isolate pre-GTP hydrolysis steps, which revealed key features that determine the relative conformational occupancy of eIF5-bound initiation complexes. To determine how allowing GTP hydrolysis impacts the conformational ensemble, we established a realtime single-molecule assay to monitor eIF5. After tethering initiation complexes, we used stopped-flow injection to add eIF5-LD655 and monitored eIF5 binding in real time (**Fig. 5A**).

**Figure 5:**
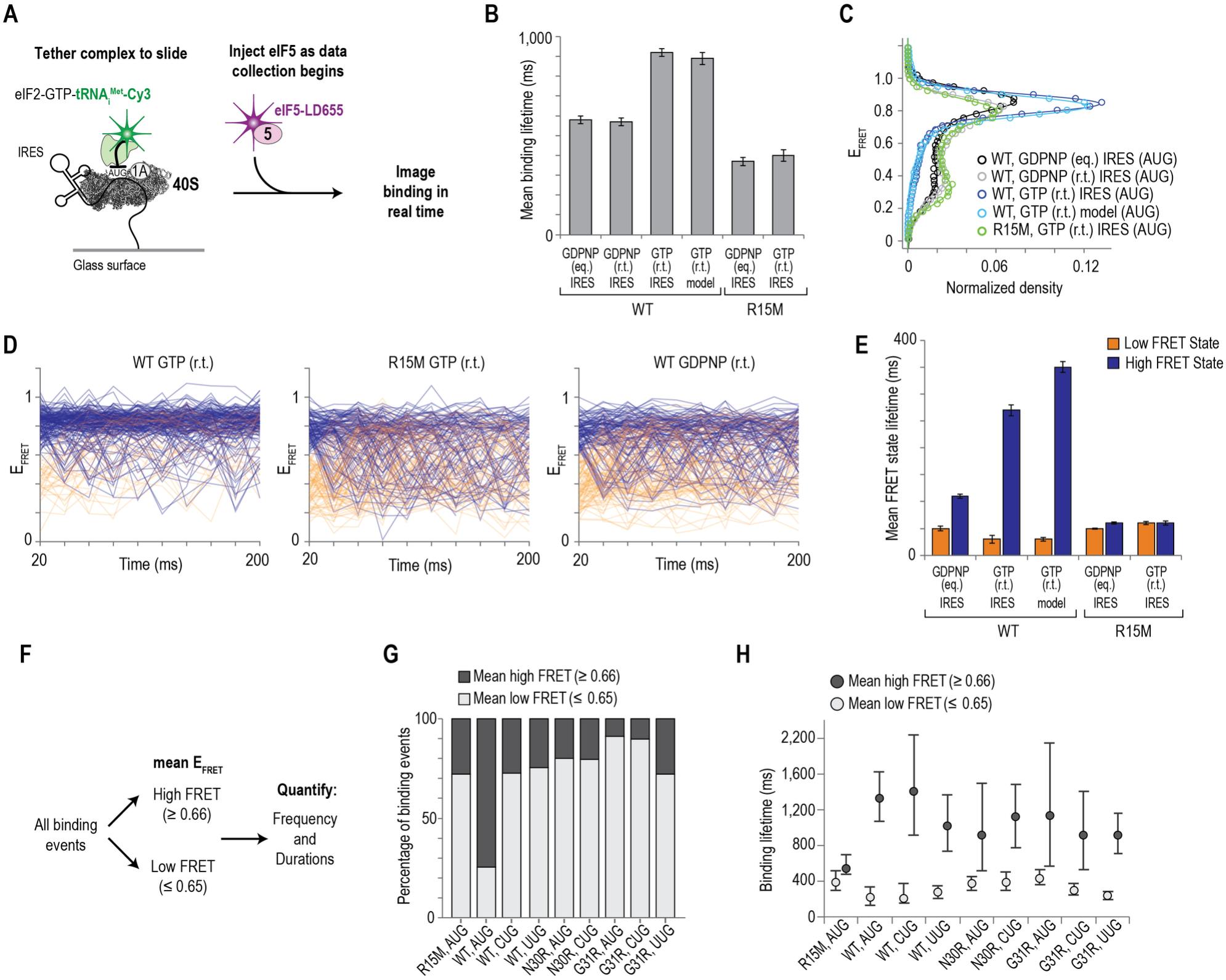
GTP hydrolysis stabilizes the high-FRET conformation. **A**, Schematic of the real-time single-molecule assay using eIF2-GTP and stopped-flow injection of eIF5-LD655. **B**, Weighted population means of wild-type eIF5 and the R15M eIF5 variant binding lifetimes in the presence of GDPNP and GTP, where ‘eq.’ refers to experiments conducted at equilibrium and ‘r.t.’ refers to experiments performed with the real-time assay. **C**, tRNA_i_-Cy3 – to – eIF5-LD655 FRET efficiency distributions observed at the indicated assay conditions; the lines represent fits to double gaussian functions. **D**, Plots of the observed FRET efficiency trajectories of two hundred binding events over the first two hundred milliseconds. Blue lines represent events that started in high FRET (≥0.66) and orange lines represent events that started in low FRET (≤ 0.65). **E**, Weighted population means of the high- and low-FRET state lifetimes for each eIF5 protein at the AUG start site in the presence of GDPNP or GTP. **F**, Schematic of data workflow to separate binding events. **G**, Plot of the percentage of binding events with a mean FRET value ≤ 0.65 (low FRET) or ≥ 0.66 (high FRET) for each eIF5 variant at the indicated start sites. **H**, Mean binding lifetimes for binding events that were on average low or high FRET, as defined in panels F and G. The error bars represent 95% C.I. of the mean yielded from bootstrapping analyses (n = 2000).

Inclusion of GTP markedly altered the conformational dynamics. eIF5 bound initiation complexes for 920 ± 20 ms and yielded a striking high-FRET population (89 %, E_FRET_ = 0.84 ± 0.08) (**Fig. 5B,C** and **Extended Data Fig. 10A,B**).

During individual binding events, eIF5 predominantly occupied the high-FRET conformation (lifetime of 270 ± 10 ms) with transient excursions into the low-FRET state (30 ± 7 ms) (**Fig. 5D,E** and **Extended Data Fig. 10C**). Similarly, eIF5 bound canonical initiation complexes assembled on a model mRNA for 890 ± 30 ms and primarily resided in the high-FRET state (lifetime of 350 ± 10 ms), with only brief transitions to the low-FRET state (30 ± 3 ms) (**Fig. 5B-E** and **Extended Data Fig. 10A, D**). In sharp contrast, prevention of GTP hydrolysis in the real-time assay – either by the hydrolysis deficient eIF5^R15M^ variant or GDPNP – shortened eIF5 binding events by 1.6–2.3-fold and reestablished rapid fluctuations between the two conformations (**Fig. 5B-E** and **Extended Data Fig. 10A-C, E**). Our findings demonstrate that GTP hydrolysis (or subsequent P_i_ release) by eIF2 stabilizes the high-FRET conformation of eIF5-bound initiation complexes, without which the complex reversibly samples both states.

We next asked whether the conformational dynamics reflect a single homogeneous class of eIF5 binding events or instead convolute distinct outcomes. Given the differential sensitivity of the two conformations to GTP hydrolysis, we classified individual binding events as either low FRET (mean E_FRET_ ≤ 0.65) or high FRET (mean E_FRET_ ≥ 0.66) (**Fig. 5F)**. For wild type eIF5 in the presence of GTP, the majority of binding events (74 %) were high FRET and had mean binding lifetime of 1,330 ms (1,070–1,630 ms, 95% C.I.) (**Fig. 5G,H** and **Supplementary Table 4**). The remaining low-FRET events had a 6-fold briefer duration of 220 ms (130–330 ms, 95% C.I.). By contrast, prevention of GTP hydrolysis by using eIF5^R15M^ reduced the fraction and duration of high-FRET binding events by 2.5-fold to 540 ms (28 %, 410–690 ms, 95% C.I.), which substantially diminished the gap in their binding lifetimes (1.3-fold difference). Thus, individual eIF5 binding events at AUG codons largely parse into two distinct conformations with different relative stabilities.

The identity of the start codon biased the distribution of eIF5 binding events in the presence of GTP. Relative to AUG, the CUG and UUG start codons reduced by nearly 3-fold the number of binding events classified as high FRET (from 74% to 28% and 25%) (**Fig. 5G** and **Extended Data Fig. 10F**). Despite this reduction, these infrequent high-FRET events on CUG and UUG codons persisted for ∼1,000–1,400 ms, essentially matching the duration on AUG (**Fig. 5H** and **Supplementary Table 4**). The predominant population of low-FRET events on CUG and UUG codons exhibited 4-7-fold briefer lifetimes, again comparable to the AUG codon (**Fig. 5H**). We observed similar, but more pronounced, effects upon substitution of N_30_ in eIF5 with arginine (**Fig. 5G,H** and **Extended Data Fig. 10G-K**). The exception was G31R, the only variant to favor the high-FRET state on UUG relative to AUG and CUG codons (**Fig. 5G** and **Extended Data Fig. 10H**), which also enhances UUG initiation in yeast^42,53^.

Allowing GTP hydrolysis by eIF2 dramatically reshapes the conformational landscape of eIF5-bound initiation complexes. With an AUG codon, the complex predominantly resides in the GTP hydrolysis-stabilized high-FRET conformation, whereas prevention of hydrolysis restores rapid interconversions between conformations. Non-AUG codons shift predominant residence to the low-FRET conformation to different extents. Importantly, AUG and non-AUG start codons differ primarily in their like-lihood of accessing the GTP hydrolysis-stabilized high-FRET conformation, which depends on the G_29_N_30_G_31_ loop of eIF5. These findings demonstrate that start codon discrimination by eIF5 occurs through biased residence in distinct conformations upstream of GTP hydrolysis (or P_i_ release), which then commits the stabilized subset of complexes to the start codon. Consistently, analogous substitutions within the G_29_N_30_G_31_ loop of yeast eIF5 enhance usage of UUG start codons^42,53^, and the G31R substitution is lethal in haploid cells^53^, linking disrupted conformational dynamics to decreased initiation fidelity.

### Destabilization of the low-FRET conformation

We next explored the molecular basis of the high- and low-FRET conformations. Using cryo-EM, we visualized the N-terminal domain of eIF5 docked adjacent to tRNA_i_ and the AUG start codon (**Fig. 1B** and **Fig. 3A**). This location agrees with prior structures^42–44^, orients N_30_ of eIF5 proximal to the codon-anticodon interface, and poises R_15_ proximal to eIF2γ, which catalyzes GTP hydrolysis (**Extended Data Fig. 8C**). It also situates the fluorescent dyes on tRNA_i_ and eIF5 to be within about 30 Å. We conclude that this position of eIF5 corresponds to the high-FRET state that becomes stabilized upon GTP hydrolysis.

To identify the structural basis of the low-FRET state, we performed additional cryo-EM analyses of eIF5-bound initiation complexes poised at an AUG or UUG start codon. After focused classifications considering only 48S particles containing eIF1A, we identified three major subpopulations of complexes (**Extended Data Fig. 1D** and **11A**). The first class contained density in the canonical eIF5 binding site and lacked density in the region vacated by the core domain of eIF2β^19,20,44,45^ (**Fig. 1A,B** and **Fig. 6A**). This class accounted for 43% of particles on the AUG codon and 10% on UUG (**Fig. 6A** and **Extended Data Fig. 11A-C**). A second class lacked density in the eIF5 binding site but contained density overlapping the eIF2β binding site. This second class dominated for the UUG codon (80%) and reduced substantially on AUG (8%). The final class did not contain density that could be attributed to eIF5. In all classes of particles, the density for tRNA_i_ closely matched the scanning-arrested state (**Extended Data Fig. 11D-F** and **12A-C**). Thus, while the tRNA_i_ conformation was similar between classes, occupancy of the eIF5 and eIF2β binding sites appeared mutually exclusive and varied with the identity of the start codon.

**Figure 6:**
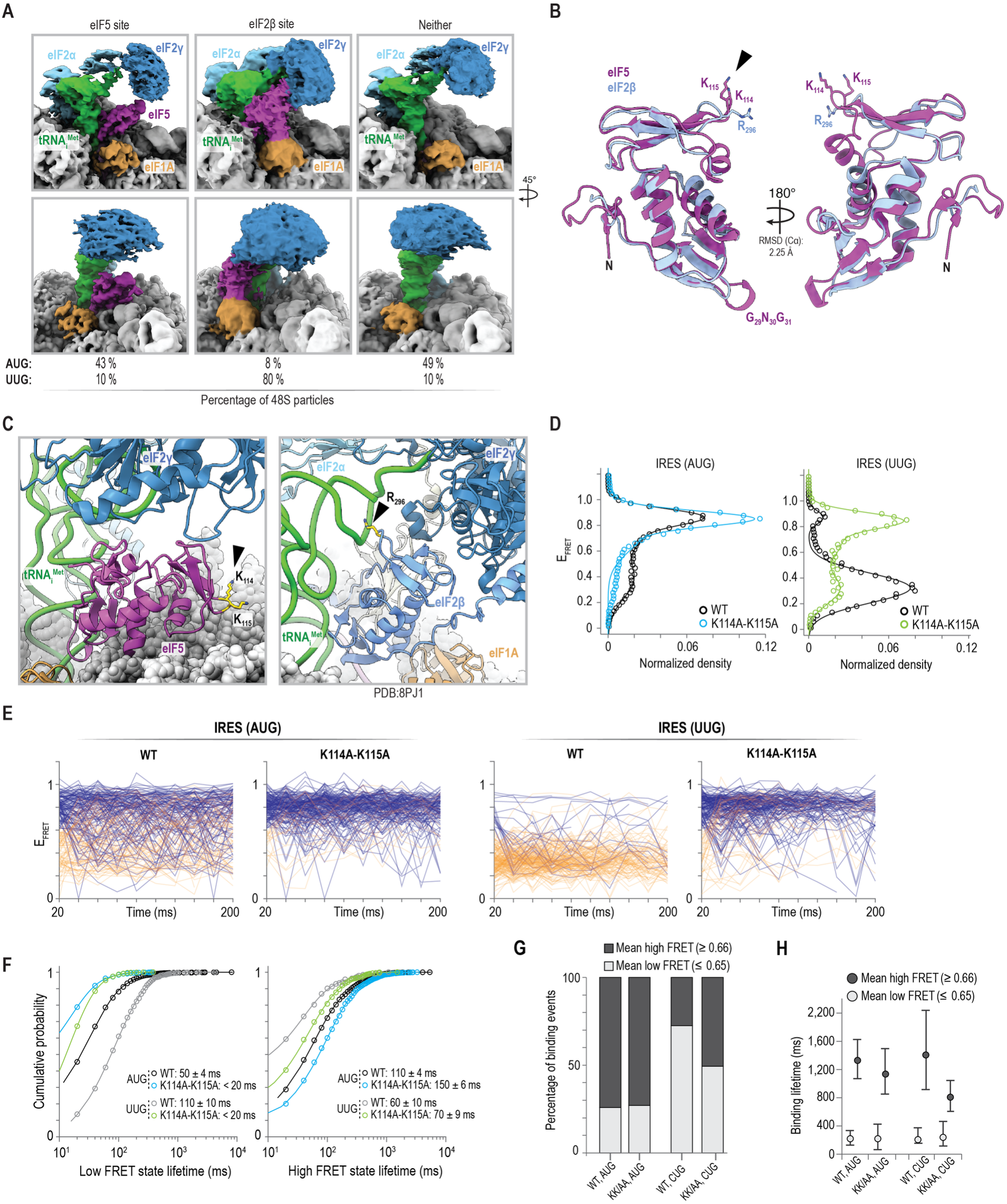
Destabilization of the low-FRET conformation. **A**, Cryo-EM density maps of three classes of 48S particles in the AUG dataset, defined by which site contained density: the canonical eIF5 binding site, the eIF2β site, or neither. Density at the eIF2β site was not assigned to a specific protein. The percent of 48S particles that belong to each class in the presence of an AUG or UUG codon is indicated. **B**, Superposition of eIF5 NTD (residues 1 – 149, purple) and eIF2β core domain (residues 198 – 314, blue). eIF5 residues K_114_K_115_ and analogous eIF2β R_296_ are shown and highlighted by the black triangle. **C**, (Left) When eIF5 resides in its canonical, high-FRET conformation, K_114_ and K_115_ are solvent exposed. (Right) Interaction of eIF2β R_296_ with tRNAi. (PDB: 8PJ1). **D**, tRNA_i_-Cy3 – to – eIF5-LD655 FRET efficiency distributions observed for wild-type eIF5 and eIF5^K114A-K115A^ with the indicated start codon; the lines represent fits to double gaussian functions. **E**, Plots of the observed FRET efficiency trajectories of two hundred binding events over the first two hundred milliseconds. Blue lines represent events that started in high FRET (≥0.66) and orange lines represent events that started in low FRET (≤ 0.65). **F**, Cumulative probability plot of the low-FRET (left) and high-FRET (right) state lifetimes for wild-type eIF5 and eIF5^K114A-K115A^ at the indicated codons. Lines represent fits to exponential functions and the derived mean times are reported on the plots. **G**, Plot of the percentage of binding events with a mean FRET value ≤ 0.65 (low FRET) or ≥ 0.66 (high FRET) for wild-type (WT) or K114A-K115A (KK/AA) eIF5 at AUG and CUG codons in the presence of GTP (real-time assay). **H**, Mean binding lifetimes for binding events that were on average low or high FRET, as defined in panel G. The error bars represent 95% C.I. of the mean yielded from bootstrapping analyses (n = 2000).

The N-terminal domain of eIF5 shares extensive structural homology with the core domain of eIF2β (2.3 Å RMSD) (**Fig. 6B**). This previously noted ancient homology^55–57^ suggested that the low-resolution density at the eIF2β position could at least partially correspond to transient repositioning of the eIF5 N-terminal domain. In this model, the eIF5 N-terminal domain and the now flexible core domain of eIF2β, anchored to eIF2γ by a conserved α-helix, dynamically occupy and exchange within the same binding site. Relative to its canonical high-FRET position (∼32 Å movement and 62° rotation), this alternative configuration of eIF5 repositions the G_29_N_30_G_31_ loop away from the codon-anticodon interface and moves R_15_ distal from the GTPase center of eIF2γ, hindering stimulation of GTP hydrolysis. It also increases the distance between the donor and acceptor dyes, consistent with the low-FRET state. To test this hypothesis, we inspected our cryo-EM models and prior structures^42–44^ to identify eIF5 residues predicted to selectively stabilize this low-FRET conformation. Analogous to R_296_ of eIF2β during scanning, residues K_114_ and K_115_ of eIF5 reside within 3 Å of U66 of tRNA_i_ (**Fig. 6B,C** and **Extended Data Fig. 12D**). These lysine residues are solvent exposed with eIF5 bound in its canonical, high-FRET position.

An eIF5 variant with substitutions to both lysine residues (K114A, K115A) dramatically reshaped the conformational landscape of eIF5-bound initiation complexes. This variant bound initiation complexes on an AUG codon for 2-fold briefer durations (280 ± 20 ms) relative to the wild-type protein (**Extended Data Fig. 12E-G**). Strikingly, the protein also almost exclusively occupied the high-FRET conformation with GDPNP present, nearly eliminating rapid interconversions between the conformations (**Fig. 6D,E**). The mean lifetime of the low-FRET conformation reduced to 20 ms on both AUG and UUG start sites (**Fig. 6F**). This value overestimates the true lifetime and indicates an at least 6-fold destabilization of the low-FRET state, as it corresponds to the temporal resolution of our experiments. The lifetime of the high-FRET conformation was modestly lengthened, likely due to missed transient excursions (<10 ms) into the low-FRET state. Substitution of K_114_ and K_115_ in eIF5 thus selectively destabilized the low-FRET conformation, which suggests that the eIF5 N-terminal domain samples the eIF2β binding site. This variant moreover decreased initiation fidelity in vitro; it increased the number of complexes that achieved the GTP-hydrolysis stabilized high-FRET conformation on a CUG codon (**Fig. 6G,H** and **Extended Data Fig. 12H**), and enhanced CUG usage in HeLa cell extracts (**Extended Data Fig. 12I**).

We last asked whether eIF5 preferentially associates with or dissociates from initiation complexes in either the high- or low-FRET conformation. The conformation first observed upon eIF5 binding in our single-molecule FRET analyses nearly always depended on features that govern its overall occupancy. The wild-type protein displayed a near equal likelihood of binding in either the high- or low-FRET conformation on an AUG codon (**Extended Data Fig. 13A-C**). Non-AUG start codons and substitutions in the G_29_N_30_G_31_ loop strongly biased initial binding toward the low-FRET state. Across nearly every eIF5 variant and start codon tested, eIF5 was more likely (1.5–2-fold) to dissociate from the complex while in the low-FRET conformation relative to the high-FRET state (**Extended Data Fig. 13D-G**). For both association and dissociation, the striking exception was the K114A-K115A double substitution in eIF5. This variant predominantly bound in the high-FRET conformation on both AUG and UUG start codons, with the latter 12-fold increased relative to the wild-type protein (**Extended Data Fig. 13C**). This variant also accelerated dissociation from the low-FRET conformation on both AUG and UUG codons (**Extended Data Fig. 13F,G**). Collectively, our single-molecule and molecular dynamics findings suggest that eIF5 rapidly decodes codonanticodon pairing, whereby proper pairing geometry or local distortions from mismatches shunt eIF5 to conformations with distinct outcomes.

## DISCUSSION

When protein synthesis begins, the translation initiation machinery must precisely establish the reading frame while also allowing regulated use of non-AUG start codons. Prior studies have largely examined how these competing needs are balanced during scanning and eIF1-dependent proofreading steps (e.g. ^20,25,26^). Here, we demonstrate that eIF5 governs a downstream conformational branchpoint once initiation complexes pause at candidate start codons (**Fig. 7**). The branchpoint depends on a strictly conserved loop in eIF5 that monitors base pairing at the codon-anticodon interface to differentially gate commitment to AUG and non-AUG codons. Our findings reframe eIF5 from a passive GTPase activating protein for eIF2 to an intrinsic regulator of start codon recognition.

**Figure 7:**
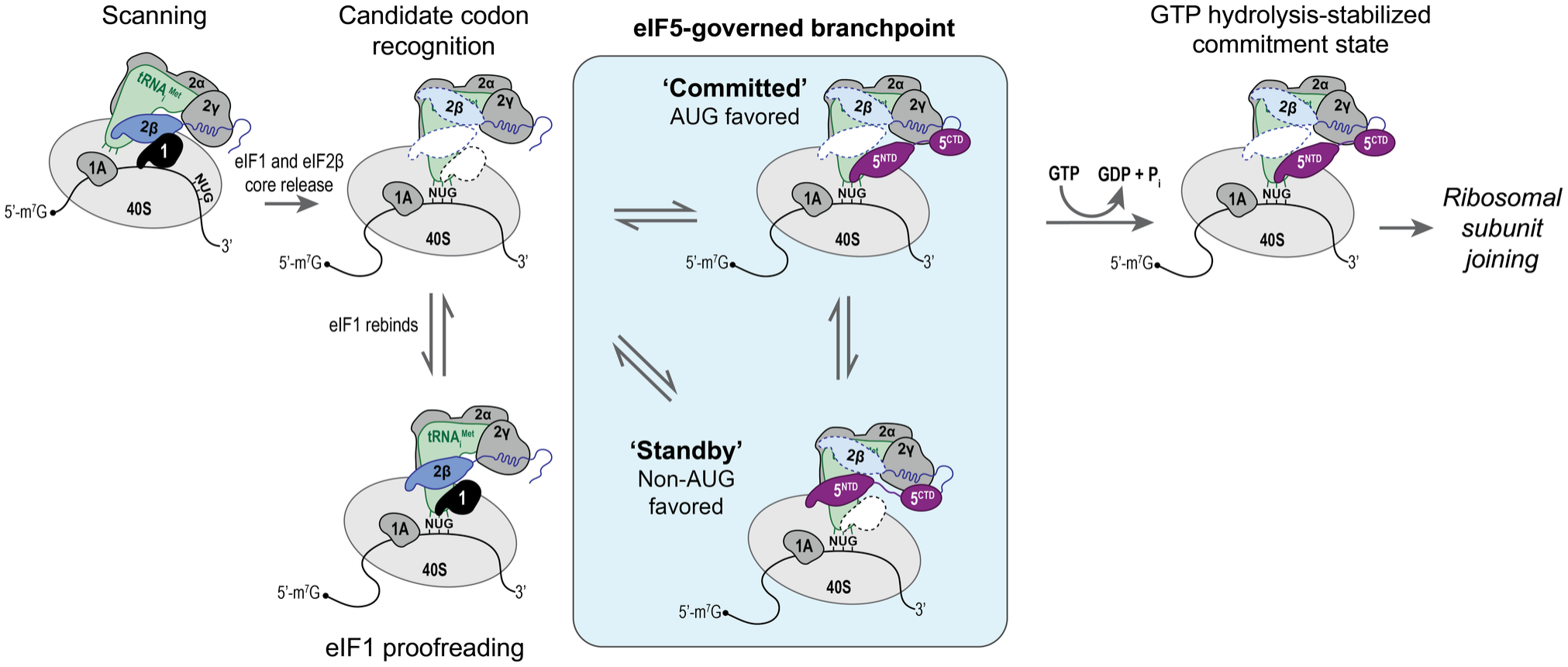
Proposed model for how eIF5 functions as a codon-sensitive branchpoint to distinguish translation start codons. Once initiation complexes pause at candidate start codons, eIF5 binds and its N-terminal domain (NTD) dynamically samples at least two conformations. The conformations correspond to distinct functional outcomes: a ‘committed’ conformation that becomes stabilized once it stimulates GTP hydrolysis by eIF2; and a ‘standby’ conformation that is insensitive to GTP hydrolysis and accelerates eIF5 release. Different start codons bias the probability that individual eIF5 binding events access the same GTP-hydrolysis stabilized conformation, intrinsically tuning usage of AUG and non-AUG codons.

We discovered that eIF5-bound initiation complexes adopt at least two distinct conformations, which we hereafter term ‘committed’ and ‘standby’. The committed (high-FRET) configuration favored by AUG codons is stabilized upon GTP hydrolysis or subsequent P_i_ release by eIF2, which ultimately allows the large ribosomal subunit to join. Non-AUG codons favor the standby (low-FRET) conformation that accelerates eIF5 departure. Prior to GTP hydrolysis, the two conformations reversibly interconvert during individual binding events on a tens-of-milliseconds timescale. Proper pairing between the start codon and anticodon of tRNA_i_ strongly biases the ensemble toward the committed conformation. Single-nucleotide substitutions bias complexes toward the standby conformation and reduce the probability that complexes achieve the committed conformation. These findings establish a key partitioning step governed by eIF5 that lies just upstream of commitment to a start codon.

The conserved G_29_N_30_G_31_ loop within eIF5 monitors codon-anticodon pairing. Our structure (**Fig. 1**) and others^42–44^ place the N-terminal domain of eIF5 within the ribosomal P site adjacent to tRNA_i_. This location corresponds to the committed conformation and overlaps the site vacated after scanning by its structural homolog and competitor, eIF1. In this position, the N_30_ residue of eIF5 resides within 3 Å of the first nucleotide of candidate start codons, allowing it to monitor the codon-anticodon interface. Indeed, alterations to G_29_N_30_G_31_ side chain chemistry, length, charge, or backbone flexibility uniformly rewire the local chemical interaction network and increase occupancy of the standby conformation, strongly reshaping the conformational ensemble. The initial conformation of the eIF5 N-terminal domain strongly correlated with canonical AUG pairing (committed) or a codon-anticodon mismatch (standby). Decoding of the codon-anticodon interface by the G_29_N_30_G_31_ loop thus occurs on a (sub-)microsecond to millisecond timescale^58^. The discrepancy in calculated hydrogen bonding lifetimes in our simulations relative to our single-molecule analyses suggests the presence of additional decoding intermediates. Regardless, our findings align with the identification of this loop as a critical determinant of start codon fidelity in a foundational yeast screen^53^ and a more recent yeast study^42^. The G_29_N_30_G_31_ loop of eIF5 thus serves as a conserved molecular sensor that couples start codon recognition to commitment.

eIF5 samples a second binding site that is likely shared with another structural homolog, eIF2β. During scanning, the core domain of eIF2β stably binds the initiation complex proximal to eIF1 and tRNA_i_, and this domain becomes dynamic once complexes pause at candidate start codons^19,20,44,45^. While other rearrangements also are possible, the standby conformation is consistent with transient residence of eIF5 in this more distal site vacated by eIF2β. This working model is supported by the correlation between our cryo-EM classifications and single-molecule FRET findings on AUG and UUG codons. The selective destabilization of the standby conformation (low-FRET state) upon disruption of eIF5 interactions predicted to be unique to the eIF2β binding site also supports this model. Such a configuration disengages the G_29_N_30_G_31_ loop from the codon-anticodon interface and moves R_15_ distal to eIF2γ, hindering GTP hydrolysis. The eIF5 N-terminal domain and the eIF2β core domain may exchange dynamically within this binding site. A dynamic exchange would be facilitated by distinct contacts between the eIF5 C-terminal domain^59,60^ and a conserved helix in eIF2β^20,46,47^ with eIF2γ, as well as interactions between the eIF5 C-terminal domain and the flexible N-terminal tail of eIF2β^60–63^, all of which may serve as anchor points. Such dynamics may explain the low resolution of the density we observed in our cryo-EM maps for the standby position and warrant future investigation. Regardless, the standby (low-FRET) configuration of the eIF5 N-terminal domain accelerates departure of the entire protein, which occurs more frequently on non-AUG codons.

Complementing eIF1-mediated proofreading, our findings establish eIF5 as a dynamic molecular switch once initiation complexes pause at candidate start codons (**Fig. 7**). During scanning, eIF1 both inhibits non-canonical pairing between mRNA codons and the tRNA_i_ anticodon. It also sterically blocks the N-terminal domain of eIF5 from binding, potentially in tandem with eIF2β, which prevents premature commitment. Once the complex halts at a candidate start codon, eIF1 departs and the core domain of eIF2β releases, allowing eIF1 and eIF5 to compete to bind. eIF1 rebinding favors codon rejection, and eIF5 binding favors commitment. Codon discrimination by eIF5 results from biased probabilities of accessing the committed conformation that becomes selectively stabilized upon GTP hydrolysis by eIF2. Rather than immediately dissociating at codon-anticodon mismatches, transient residence of eIF5 in a standby configuration may allow eIF5 repeated attempts to commit complexes to non-AUG codons. eIF5 variants that prolong residence in the standby conformation indeed enhance relative usage of non-AUG codons in yeast^42,53^. The standby state nonetheless accelerates eIF5 dissociation, which occurs more frequently on non-AUG codons and restarts its competition with eIF1. This conformational branchpoint controlled by eIF5 thus provides a mechanism for how translation initiation achieves high specificity for AUG codons while allowing regulated use of non-AUG codons and suggests potential functions for conserved structural homology between initiation factors.

## ACKNOWLEDGEMENTS

We are grateful to members of the Lapointe, Campbell, and Subramaniam labs for helpful guidance, discussions, and feedback. We appreciate helpful feedback from T. Dever, M. Grimes, and M. Lawson on the manuscript, advice from R. Grosely and on the extract assays, and discussions with N. Fischer on the findings. This work was funded, in part, by the National Institutes of Health (R35GM160398 to C.P.L.; R35GM119835 to A.R.S; R35GM147414 to M.G.C.; T32CA080416 to M.C.C.; T32GM153507 to T.C.C.; F32GM163369 to R.R.C.), the Rita Allen Foundation (Scholar Award to C.P.L.), the Damon Runyon Cancer Research Foundation (DFS-49-22 to C.P.L.), the Pew Charitable Trusts (Biomedical Scholar Award to M.G.C), and the Mahan Fellowship (to M.C.C., made possible by funding from M. Mahan and N. Mahan). This research also was supported by the Proteomics and Metabolomics Shared Resource, RRID:SCR_022618, of the Fred Hutch/University of Washington/Seattle Children’s Cancer Consortium (P30CA015704) and the Fred Hutch Scientific Computing, NIH grants S10-OD-020069 and S10-OD-028685. Electron microscopy data were generated using the Fred Hutchinson Cancer Center Electron Microscopy shared resource, supported in part by the Cancer Center Support Grant P30CA015704-40.

## AUTHOR CONTRIBUTIONS STATEMENT

These authors contributed equally: SFM and MCC. Conceptualization: SFM, MCC, CPL Methodology: SFM, MCC, TCC, CPL Investigation: SFM, MCC, TCC, RRC, NP, EMA, VMS, MGC, CPL Visualization: SFM, MCC, TCC, CPL Funding acquisition: ARS, MGC, CPL Project administration: MGC, CPL Supervision: MGC, CPL Writing – original draft: SFM, CPL Writing – review & editing: SFM, MCC, TCC, RRC, NP, EMA, VMS, ARS, MGC, CPL

## DATA AVAILABILITY

All single-molecule traces and source data needed to recapitulate single-molecule figure plots throughout the manuscript are available for download from GitHub: https://github.com/LapointeLab/eIF5-2026-paper. Specific requests can be made using the Issue feature or by email request to C.P.L. Structure images were generated using published models (PDB IDs: 8PJ1, 8PJ2) and ChimeraX-1.71.1 software. Atomic models and cryoEM maps of the initiation complex on AUG are deposited to the Protein Data Bank (PDB: 11NV) and to the Electron Microscopy Data Bank (EMD: 75872). Density maps of initiation complexes in other eIF5 states are deposited to the Electron Microscopy Data bank (EMD: 76218 (AUG-eIF5 absent), 76219 (AUG-eIF5 standby), 76220 (UUG-eIF5 absent), 76221 (UUG-eIF5 committed), 76222 (UUG-eIF5 standby). Molecular dynamics topology, initial and final coordinates are available on GitHub: https://github.com/LapointeLab/eIF5-2026-paper. Source data are provided with the manuscript.

## CODE AVAILABILITY

Related analysis code is publicly available on GitHub: https://github.com/LapointeLab/eIF5-2026-paper.

## METHODS

### Labeled eIF5

Purification and fluorescent labeling of functional eIF5 was performed as described^26^ using the following protocol. Synthetic DNA that encoded human eIF5 with an N-terminal ybbR tag was purchased from IDT and inserted into the pET28 backbone, which contained a TEV protease cleavage site between the 6×His tag and the ybbR tag. Mutations in eIF5 were introduced using site-directed mutagenesis, which were verified by sequencing. Rosetta2 cells transformed with the expression plas-mid were grown to an OD of 0.5 and induced with 0.5 mM IPTG for 4 h at 30 °C. Cells were harvested by centrifugation at 5000 RPM for 10 min at 4°C in a F9-6x1000 LEX rotor (Thermo Scientific) and the remaining cell pellets were frozen at -20°C. Frozen cell pellets were thawed and resuspended in lysis buffer (20 mM Tris-HCl pH 8.0, 300 mM NaCl, 10% (v/v) glycerol, 20 mM imidazole, 10 µM zinc sulfate, and 5 mM β-mercaptoethanol) supplemented with protease inhibitors. Cells were lysed by sonication in lysis buffer and lysates were cleared by centrifugation at 16,000 RPM for 45 min at 4°C in a F20-12×50 LEX rotor (Thermo Scientific) followed by filtration through a 0.22-µm syringe filter. Clarified lysate was loaded onto a Ni-NTA gravity flow column equilibrated in lysis buffer, washed with 10 column volumes (CVs) of lysis buffer, 10 CVs of wash buffer (20 mM Tris-HCl pH 8.0, 800 mM NaCl, 10% (v/v) glycerol, 40 mM imidazole, 10 µM zinc sulfate, and 5 mM β-mercaptoethanol) and 10 CVs of lysis buffer. Recombinant proteins were eluted with five sequential CVs of elution buffer (20 mM Tris-HCl pH 8.0, 300 mM NaCl, 10% (v/v) glycerol, 300 mM imidazole, 10 µM zinc sulfate, and 5 mM β-mercaptoethanol). Fractions with recombinant protein were identified by SDS–PAGE analysis. The relevant fractions were dialyzed overnight at 4 °C into tobacco etch virus (TEV) cleavage buffer (20 mM Tris-HCl pH 7.5, 200 mM NaCl, 10% (v/v) glycerol, 10 µM zinc sulfate, and 1 mM DTT) in the presence of excess TEV protease. 30mM imidazole was added to the dialyzed protein sample and the TEV protease and the cleaved 6×His–MBP tag were removed using a subtractive Ni-NTA gravity column equilibrated in TEV buffer supplemented with 27 mM imidazole with the flowthrough collected. The protein sample was diluted to 75 mM NaCl, applied to a 1-ml Q FF ion-exchange column and eluted using a gradient of 50 mM to 1M NaCl in the absence of reducing agents. Fractions that contained eIF5 at >95% purity were concentrated to ∼150 μM, frozen in liquid N2 and stored at ™80 °C. To fluorescently label the purified protein, ∼30 μM ybbR–eIF5 was supplemented with 10 mM MgCl_2_ and 1 mM DTT and incubated at 37 °C for 60 min in the presence of 4 μM sfp synthase enzyme and 60 μM Cy5–CoA or LD655–CoA substrate. Free dye was removed by purification over 10DG-desalting columns (Bio-Rad, 7322010) equilibrated in SEC buffer (20 mM HEPES-KOH pH 7.5, 200 mM KOAc, 10% (v/v) glycerol, 10 µM zinc sulfate, 1 mM DTT). The labeled protein was supplemented with 30 mM imidazole and sfp synthase was removed using a second subtractive Ni-NTA gravity column equilibrated in SEC buffer supplemented with 30 mM imidazole, with the flow through collected. The labeled protein then was purified using a 23-ml SD75 size-exclusion chromatography column. Fractions that contained ybbR–eIF5 at >95% purity were concentrated to ∼10 μM (by dye; ∼50% labeled), frozen in liquid N2 and stored at ™80 °C.

### Labeled initiator tRNA

HPLC purified, synthetic human Met-tRNA_i_ without native modifications and with U46 replaced with Uridine-C6 Amino linker conjugated to Cyanine 3 SE was purchased from TriLink. Prior to aminoacylation, the fluorescently-labeled tRNA_i_-Cy3 was purified using phenol:chloroform extraction and ethanol precipitation. Aminoacylation with methionine was performed as described^26,64^. The charging efficiency was > 90% based on acid urea PAGE analysis.

### Unlabeled eIFs and ribosomal subunits

Recombinant eIF1^65^, eIF1A^65^, eIF4E^66,67^, and eIF4AI^66^ proteins were purified as described. Native human eIF2 and eIF3 were purified as described^65,68^. Native human 40S and 60S ribosomal subunits were purified from either HEK293T cell line as described^69,70^. Cells were not tested for mycoplasma contamination.

eIF4GI (amino acids 557-1599) was expressed and purified as described with some modifications^71^. Briefly, eIF4G was expressed as a MBP-6x-His fusion protein, with a TEV cleavage site between the 6x-His tag and eIF4G. Briefly, NEBExpress cells were transformed with the expression plasmid and grown to an OD_600_ of ∼0.6 at 37 ºC, cooled to 18 ºC, and protein expression was induced with 0.5 mM IPTG for 16 hours. Cell pellets were resuspended in lysis buffer (20 mM Tris pH 8, 300 mM NaCl, 10% glycerol, 15 mM Imidazole, 5 mM βME, 0.1% Triton X-100) supplemented with Pierce Protease Inhibitor Tablets (Thermofisher), lysed via sonication, and clarified via centrifugation. Lysate was run over an Ni-NTA gravity column equilibrated in lysis buffer, and 6x-His-MBP-eIF4G was eluted in lysis buffer supplemented with 300 mM imidazole. Cleavage of the MBP tag was carried out overnight in the presence of excess TEV, and cleaved 6x-His-MBP was subsequently removed using an Ni-NTA gravity column. eIF4G was further purified using a HiTrap Heparin HP 1 mL column (Cytiva) eluted over a gradient from 100 to 500 mM Potassium Acetate, and fractions containing eIF4G were further purified using a HiLoad 16/600 Superdex200 column in SEC buffer containing 20 mM HEPES pH 7.3, 250 mM Potassium Acetate, 10% glycerol, 1 mM DTT. Fractions containing eIF4G were concentrated and stored at -80 ºC.

eIF4B was purified as described with some modifications^72,73^. Rosetta2 cells were transformed with 6x-His tagged eIF4B expression plasmid with TEV cleavage site between the 6x-His tag and eIF4B. Cells were grown at 37 ºC to an OD_600_ of ∼0.6, cooled to 18 ºC, and induced with 1 mM IPTG for 16 hours. Cells were resuspended in lysis buffer (20 mM Tris pH 8, 400 mM KCl, 10% glycerol, 30 mM Imidazole, 10 mM βME) supplemented with Pierce Protease Inhibitor Tablets (Thermofisher), lysed via sonication, and clarified via centrifugation. Lysate was run over an Ni-NTA gravity column equilibrated in lysis buffer and 6x-His-eIF4B was eluted with 250 mM imidazole. The 6x-His tag was cleaved using excess TEV overnight, and cleaved 6x-His and TEV were removed using an Ni-NTA gravity column. eIF4B was further purified using a HiTrap Heparin HP 1 mL column (Cytiva) eluted over a gradient from 125 to 550 mM KCl. Fractions containing eIF4B were further purified using a Superdex 200 Increase 10/300 GL column in Sizing Buffer (20 mM HEPES pH 7.3, 300 mM Potassium Acetate, 10% glycerol, 1 mM DTT). Fractions containing eIF4B were concentrated and stored at -80 ºC.

### IRES RNAs

HCV IRES RNAs were prepared as described previously^69^. A synthetic DNA was purchased from IDT that contained a 5’ flanking sequence (pUC19 backbone sequence), a T7 promoter (TAATACGACTCACTATAG), and the HCV IRES. For HCV IRES(ΔdII) constructs, IRES domain II was replaced with sequence (GGACTTCGGTCC) that encodes a stem and tetraloop to promote folding of IRES domain III^74^. In all constructs, downstream of the AUG encoded the rest of domain IV, the HCV core sequence, and an artificial 3’ extension. The templates were generated via standard PCRs using NEB Phusion polymerase (25 cycles). Purified PCR products were used as the templates for in vitro transcription with T7 MEGAScript T7 Transcription kit (ThermoFisher, #AMB13345). Standard reaction conditions were used, which was followed by treatment with Turbo DNase. The RNAs were purified using the Gene-JET RNA Purification Kit (ThermoFisher, #K0732). The 3’-terminus was biotinylated by potassium periodate oxidation followed by reaction with biotin-hydrazide, as described ^52^. The full sequences are available in **Supplementary Table 5**.

### Unstructured model mRNAs

As described previously^26^, DNA that encoded the reverse complement of the RNA and T7 promoter sequences were purchased from IDT as single-stranded oligos. After hybridizing a second oligo that encoded the sense T7 promoter sequence, the RNAs were in vitro transcribed using T7 RNA polymerase for 3 hrs at 37 °C and treated with Turbo DNase. The transcribed RNAs were capped with m^7^G using the one-step NEB faustovirus capping and 2’O-methylathion protocol. The 3’-terminus was biotinylated by potassium periodate oxidation followed by reaction with biotin-hydrazide, as described ^52^. At all steps, purifications were done using standard phenol:chloroform extractions and ethanol precipitations. The full sequence is available in **Supplementary Table 5**. As indicated, different start sites were inserted in place of the indicated AUG codon.

### Single-molecule surface preparation and passivation

Glass slides (Fisherbrand, #125444) and glass coverslips (VWR, #48404-467) were cleaned and passivated with a PEG/PEG-biotin mixture based on a previously described protocol^75^. Briefly, slides and coverslips were washed for 15 min by sonication immersed sequentially in water,100 % ethanol, water, and 100 % ethanol. Following each wash, they were rinsed five times in water. Slides and coverslips were then sonicated immersed in 1 M NaOH for 15 minutes, rinsed five times using water, dipped in acetone (Avantor J.T. Baker), and then allowed to air dry. The cleaned slides and coverslips were treated with a 1.75 % Vectabond solution (Vector Laboratories) diluted in acetone (v/v) for 3 min, rinsed using a 50 % water-acetone solution (v/v), dipped in 100 % acetone, and allowed to air dry at room temperature. The dried slide and cover slip surfaces were treated for 2-3 hours with a 1:6 mixture of biotin-PEG-5k-SVA and mPEG-5k-SVA (Laysan Bio, BIO-SVA-5k; M-SVA-5k) resuspended in ice cold 100 mM potassium borate (pH 8.4). Treated slides and coverslips were rinsed five times using water and stored under vacuum until use.

Immediately prior to imaging, slides and glass coverslips were assembled using double-sided tape to generate separate imaging channels, with the sides sealed using epoxy. The channels were washed using TP50 buffer (50mM Tris-acetate pH 7.5, 100 mM potassium acetate). Washed coverslips were coated with neutravidin using a 5-min incubation with 1 µM neutravidin diluted in TP50 buffer supplemented with 0.7 mg ml™1 UltraPure BSA and 1.3 μM of preannealed DNA blocking oligos (CGTTTACACGTGGGGTCC CAAGCACGCGGCTACTA-GATCACGGCTCAGCT and AGCTGAGCCGT GATCTAG-TAGCCGCGTGCTTGGGACCCCACGTGTAAACG). The imaging surface was washed again with TP50 buffer and remained immersed in TP50 until complexes were tethered.

### Initiation complex assembly

#### RNA refolding

All IRES RNAs were diluted to 600 nM in refolding buffer (20 mM Hepes-KOH pH 7.3, 1 mM EDTA pH 8, and 70 mM potassium acetate) and heated to 95 ºC for 2 min. The IRES was then slow cooled to room temperature over 20-30 minutes, and 4 mM magnesium acetate was added to quench the EDTA. The refolded IRES was kept on ice until use. The unstructured model mRNAs were refolded using an identical protocol, except the RNAs were placed immediately on ice after the 2 min incubation at 95 ºC.

#### Initiation reaction and imaging buffers

In all experiments, the initiation reaction buffer was: 20 mM Hepes-KOH pH 7.3, 70 mM potassium acetate, 2.5 mM magnesium acetate, 0.25 mM spermidine, and 0.2 mg/mL creatine phos-phokinase. In all experiments, the imaging buffer was the initiation buffer supplemented with casein (100 μg ml^™1^), 2.5 mM TSY, and an oxygen-scavenging system: 2 mM protocatechuic acid and 50 nM protocatechuate-3,4-dioxygenase^76^.

#### eIF2-tRNA_i_ complexes

We incubated 3.3 µM eIF2, 1 mM GDPNP (or 1 mM GTP when noted), and 1 mM magnesium acetate in initiation buffer supplemented with an additional 0.3 mg/mL creatine phosphokinase for 10 min at 37 ºC. This step promoted saturation of purified eIF2 with GDPNP (or GTP). We then added 3 µM of Met-tRNA_i_^Met^-Cy3 and incubated for an additional 5 min at 37 ºC.

#### Initiation complex formation

To form IRES-bound initiation complexes prior to tethering, we incubated 200 nM 40S subunits, 400 nM eIF1A, 100 nM refolded IRES, and 400 nM eIF2-tRNA_i_-Cy3 complex for 15 min at 37 ºC in initiation buffer. To form canonical complexes on the unstructured model mRNA, we incubated 200 nM 40S subunits, 400 nM eIF1A, 400 nM eIF1, 400 nM eIF3, 400 nM eIF2-tRNA_i_-Cy3 complex, and 100 nM model unstructured model RNA for 15 min at 37 ºC in initiation buffer. All complexes were stored on ice until use.

### Single-molecule microscope

All measurements were performed using objective-based total internal reflection fluorescence (TIRF) on a Nikon Eclipse Ti2-E inverted microscope. The microscope contained an OBIS 532 nm LS 150 mW excitation laser and a Nikon Apochromat 100X TIRF oil immersion objective (1.49 NA). To clean and separate the excitation and emission paths, we used two multi-band pass filters (excitation: ZET405/488/532/642 X; emission: ZET405/488/532/642 M) in tandem with a multi-band dichroic beamsplitter (ZT488/532/633/830/1064rpc, UF2). The emission path was separated further into Cy3 and Cy5/LD655 fluorescence bands using a Cairn Multisplit V2 (1X magnification) equipped with a single filter/dichroic cube in position 1. The cube contained a T635lpxr-UF2 dichroic beamsplitter and ET585/65m (Cy3) and ET685/70m (Cy5/LD655) emission filters. All filters and dichroic beamsplitters were purchased from Chroma Technology Corp. Photons in each band were detected using a Teledyne Photometrics Kinetix 10 MP sCMOS camera (6.5 µm x 6.5 µm pixels, 3200 x 3200 pixels). Fluorescence intensities at a resolution of 100 ms exposures (10 frames per second) were collected using 16-bit Sub-electron mode, and data with 20 ms exposures collected using 12-bit sensitivity mode. All data were collected using perfect focus and without binning pixels. To enable nearly real-time sample delivery, the microscope integrates a KDS Legato 110 syringe pump. The microscope, excitation laser, camera, and syringe pump were all operated using the Nikon Elements software suite (version 6.02.03).

### Single-molecule data collection

#### Equilibrium imaging

In all equilibrium experiments, we used eIF2-GDPNP-tRNA_i_-Cy3 complexes to facilitate analysis of multiple eIF5 binding events per initiation complex and isolate pre-GTP hydrolysis effects. The relevant initiation complex was tethered to the neutravidincoated imaging surface for 5 min at room temperature. The surface was washed once with 30 µL of initiation buffer. For imaging of IRES complexes, we immersed complexes in imaging buffer supplemented with 50 nM eIF1A and eIF5-Cy5 or eIF5-LD655 (typically at 30 nM concentration by dye; 50-70% labeling efficiency). For imaging canonical complexes, we immersed complexes in imaging buffer supplemented with 50 nM eIF1A, 50 nM eIF1, and eIF5-Cy5 or eIF5-LD655.

#### Real time imaging

The relevant initiation complex was tethered to the neutravidin-coated imaging surface as above. Following one wash using initiation buffer, the tethered complexes were immersed in imaging buffer (lacking eIF5-LD655, but containing unlabeled eIF1A and/or eIF1, as noted above), the density of tethered complexes was briefly verified, and the focus was finely adjusted. With the excitation laser off, a new field of view was selected, and 20 µL of imaging buffer supplemented with 30 nM eIF5-LD655 and 50 nM of eIF1A and/or eIF1 was injected using the attached syringe pump. Following a 10 ms delay to allow surface relaxation, fluorescence data were collected at 20 ms temporal resolution.

#### Full initiation imaging

To form the 43S pre-initiation complex, we incubated 200 nM 40S subunits, 400 nM eIF1A, 400 nM eIF1, 400 nM eIF3, and 400 nM eIF2-GDPNP-tRNA_i_-Cy3 complex for 15 min at 37 ºC in initiation buffer. Separately, the model mRNA was tethered to the neutravidin-coated imaging surface as above. Following one wash using initiation buffer, the tethered mRNA was immersed in imaging buffer supplemented with 1 µM eIF4A, 200 nM eIF4E, 200 nM eIF4G, 200 nM eIF4B, 1 mM ATP, and 1 mM magnesium acetate, and the focus was finely adjusted. With the excitation laser off, a new field of view was selected, and 20 µL of imaging buffer supplemented with 30 nM eIF5-LD655, 6 nM 43S complex including eIF2-GDPNP-tRNA_i_-Cy3, 100 nM eIF1A, 100 nM eIF1, 1 µM eIF4A, 1 mM ATP, and 1 mM magnesium acetate was injected using the attached syringe pump. Following a 10 ms delay to allow surface relaxation, fluorescence data were collected at 20 ms temporal resolution.

### Single-molecule data analyses

Prior to each data collection session, we collected 50 frame movies at 100 ms exposures of 0.1 µm Tetraspeck beads (ThermoFisher, #T7279) to ensure robust pixel registration across the Cy3 and Cy5/LD655 fluorescence channels. All raw data of fluorescence intensities over time were converted from the proprietary ND2 format to multipage .tif stacks using Nikon Elements software. All subsequent data processing steps were performed using MATLAB R2023-b functions or modules operated within the MATLAB environment. Cy3 and Cy5/LD655 fluorescence intensities were extracted from single spots and automatically background corrected using SPARTAN 3.7.0^77^ and the appropriate pixel registration matrix. Single complexes were indicated by the presence of single-step photobleaching of the Cy3 donor signal. FRET on and off states were assigned automatically using vbFRET (version 10)^78^, which were visually inspected and manually corrected as needed.

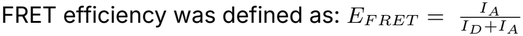

where *I*D and *I*A represent fluorescence intensities of the donor (Cy3) and acceptor (Cy5 or LD655) fluorophores. FRET efficiency distributions were determined by aggregating all observed FRET efficiencies in each frame during all binding events, which then were binned (50 bins, ™0.2 to 1.2), counted, and normalized to the total number of frames (i.e., efficiency values) in each dataset. Since only frames observed during eIF5 binding events were included, the distributions approach zero at E_FRET_ = 0. Distributions were fit to gaussian functions to determine the reported means and standard deviations of each population.

To determine eIF5 binding kinetics, we used the manually corrected Cy3-to-Cy5/LD655 FRET-on and FRET-off states identified using vbFRET to demarcate eIF5 binding events. For association kinetics (i.e., rebinding times), we quantified the time elapsed between eIF5 binding events on individual initiation complexes; that is, the time elapsed from loss of an eIF5 FRET signal to appearance of the next eIF5 FRET event on single complexes. The observed times at the indicated eIF5 concentrations were converted into cumulative probability functions (cdfcalc, MATLAB) and fit with exponential functions to derive the association rate constants. Linear regression of the observed rate constant at each concentration was used to determine the reported association rate. For dissociation kinetics (i.e., binding lifetimes), we quantified the duration of each eIF5 FRET event, converted the observed times to cumulative probability functions, and fit the data with exponential functions to derive the dissociation rates (i.e., mean lifetimes or durations). For all kinetics analyses, the base exponential function was defined as:

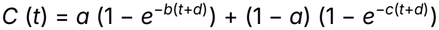

where *t* is time (in s), *b* and *c* are rates, and *d* is an adjustment factor.

To generate FRET Efficiency density plots (**Figs. 2B, 2H**), we counted the frequency of the observed FRET efficiency in each frame (using 50 bins), generated 150 contour levels, and plotted the contours across the first 500 ms of all observed eIF5 binding events in the experiment. Time 0 was defined as the start of the binding event.

To generate FRET trajectory plots, we overlaid the observed FRET efficiencies from the first 200 ms of 200 individual eIF5 binding events. In these analyses, the first frame of each binding event was excluded because eIF5 could have bound at any point during the 20 ms exposure for that frame, which convolutes the observed FRET efficiency in that frame. We color-coded binding events according to their initial FRET efficiency, with blue denoting events that begin in high FRET (≥0.66) and orange for low FRET (≤0.65). This cutoff represents the minimum between low- and high-FRET populations in the aggregate FRET efficiency distribution (**Extended Data Fig. 3G**).

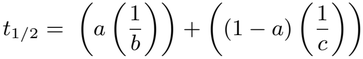

To examine FRET states and transitions, we uniformly applied a threshold-based definition of low- and high-FRET states to 20 ms exposure data. In equilibrium experiments with an AUG start codon, the aggregate FRET efficiency distribution was modeled well by three populations: a major high-FRET population (0.86 ± 0.08, 48%), a high-FRET shoulder (0.75 ± 0.15, 30%), and a minor low-FRET population (0.41 ± 0.21, 22%) (**Extended Data Fig. 3G**). There was a minimum between the low-FRET and overlapping high-FRET populations, centered at ∼0.65. Based on this distribution, we defined the low-FRET state as FRET efficiencies ≤ 0.65 and the high-FRET state as efficiencies ≥ 0.66, which we applied uniformly across all datasets. During individual binding events, the Cy3 donor and LD655 acceptor fluorescence intensities detected in each frame were strongly anti-correlated (median Pearson r = -0.8 to -0.86 across replicates), indicative of rapid and well-resolved FRET fluctuations at our 20 ms timescale. Transitions between the FRET states were identified by transits through the FRET efficiency threshold (0.65) during individual binding events, and highly similar results were obtained when using thresholds of 0.55 and 0.6. The observed durations (i.e., lifetimes) of the low- and high-FRET states were modeled by cumulative probability and double-exponential functions to derive the reported population-weighted mean lifetimes. This approach estimates the stability of each state and facilitates comparisons of relative stabilities across experiments without a predefined kinetic model. The reported FRET state life-times thus should be interpreted as upper bounds of the durations.

### Association and Dissociation conformation analyses

We used vbFRET and manual corrections to define the frame where an eIF5 binding event begins and the frame where the binding event ends. Each eIF5 molecule may bind at any point during the 20 ms exposure of frame 1: after 0 ms, 2 ms, 5 ms, 15 ms, etc. The same effect is true for dissociation: eIF5 may have departed at any point during the 20 ms exposure window of the final frame. This means that the first frame (frame 1) and final frame of each eIF5 binding event provide a time censored FRET efficiency value, which artificially biases the observed FRET to a lower value.

#### Association analyses

To examine any potential conformational preference as eIF5 binds (associates), we calculated the instances of observed high-FRET relative to all detected frames for frame 1, 2, 3, 4, and 5. To estimate the error, we bootstrapped the observed percentage (n = 2000) using a per molecule resampling strategy to calculate 95 % C.I., since events on individual complexes are not independent. We compared the observed percent high FRET to the fraction of all frames with the observed FRET efficiency detected across all binding events. Consistent with time censoring of frame 1, we detected a large increase in high-FRET association between frames 1 and 2. The observed percentage plateaued from frames 2-5 at ∼55%, nearing the fraction of all frames detected in high FRET (aggregate). Given this observation and since the mean duration of eIF5 residence in both the low- and high-FRET states is longer than 20 ms, we analyzed frame 2 as a proxy for detecting whether eIF5 bound preferentially in either low- or high-FRET states.

#### Dissociation analyses

To uncover whether eIF5 dissociated preferentially from the low- or high-FRET state, we considered at least three possibilities as a binding event progressed from one frame to another: 1) the FRET Efficiency (i.e. state) is maintained; 2) the FRET state switches from high-to low-FRET, or from low-to high-FRET; or 3) the binding event ends with eIF5 dissociation. Our reported low-FRET and high-FRET state lifetimes convolute outcomes 2 and 3 (state switch or dissociation). We therefore calculated the ‘dissociation probability’ of the indicated frames. For both low FRET and high FRET, we determined the number of observed instances in a given frame that led to dissociation, and we calculated the ratio of the observed instances relative to the total number of frames detected with that FRET state across the full duration of all binding events. We also compared the ratio of the low-FRET and high-FRET dissociation probabilities of given frames. The reported errors represent 95 % C.I. determined using bootstrapping (n = 2000) and a per molecule resampling strategy, since events on individual complexes are not independent. We compared the dissociation probabilities and dissociation ratios of the final, 2nd to last, 3rd to last, 4th to last, and 5th to last frames of the binding event.

### nanoLuciferase mRNA generation

Geneblocks were purchased from IDT that contained a T7 promoter and nLuc coding sequence (Promega) flanked by 5’ and 3’ UTRs derived from ß-globin transcript as described^32,69^. DNA templates were generated via standard PCRs using NEB Phusion polymerase (25 cycles). Purified PCR products were used as the templates for in vitro transcription with T7 MEGAScript T7 Transcription kit (ThermoFisher, #AMB13345). Standard reaction conditions were used, which was followed by treatment with Turbo DNase. The RNAs were purified using the GeneJET RNA Purification Kit (ThermoFisher, #K0732). The transcribed RNAs were capped with m^7^G using the one-step NEB faustovirus capping and 2’-O-methylathion protocol. Capped mRNAs were purified using GeneJET RNA Purification Kit (ThermoFisher, #K0732) and analyzed on a 1.5% agarose gel.

### In vitro extract-based translation assays

Standard HeLa cell-free translation (ThermoFisher, #88882) reactions (10 µL) were programmed with a final mRNA concentration of 100 nM and 5 µM of the indicated eIF5 protein. An equal volume of eIF5 SEC (storage) buffer (without protein) was added to a control reaction to account for buffer effects on IVT activity. IVT reactions were incubated at 37 °C for 45 min and then immediately transferred to ice and diluted 1:1 with cold Nano-Glo Buffer (Promega, #N1120). An equal volume of 1:50 diluted Nano-Glo substrate was added to each reaction. Samples (90% of total volume) were loaded into non-adjacent wells of a 384-well plate. Sample luminescence was measured 11 min post nGlow solution addition using a LUMIstar Omega plate reader (25 °C, gain of 3600). Prism8 (Graphpad 10.5.0) was used for all analyses.

### Cryo-EM sample preparation

### Initiation complex formation

To form IRES-bound initiation complexes, we incubated 300 nM 40S subunits, 2µM eIF1A, 500 nM refolded IRES with the canonical AUG start codon, and 1 µM eIF2-GDPNP-tRNA_i_-Cy3 complex for 10 min at 37 ºC in initiation buffer. We then added 1.5 µM of wild-type eIF5-LD655 and incubated the complex for 15 min at 37 ºC.

#### BS3 crosslinking

Initiation complexes were cross-linked with BS3 to increase complex stability for cryo-EM^44,45^. 2 mg of BS3 (bis(sulfosuccinimidyl)suberate) (Thermo Scientific, #A39266) was resuspended to 10 mM in 20 mM Hepes-KOH pH 7.3, 70 mM potassium acetate, 2.5 mM magnesium acetate. We incubated 250 nM 48S complex with eIF5 (based on 40S subunit concentration) and 1.5 mM BS3 on ice for 45 min.

The assembled initiation complex (3 µl) was applied to Quantifoil R 1.2/1.3, UT, 300-mesh Cu grids that were glow discharged (15 sec, 15 mA). Grids were blotted using a Vitrobot Mark IV (Thermo Fisher Scientific) and a blot time of 4, blot force of 1 to 2, at 4C and 100% humidity. Grids were immediately vitrified in liquid ethane cooled by liquid nitrogen and stored in liquid nitrogen until use.

### Cryo-EM data collection and processing

Data were collected on a 200 kV Glacios cryo-transmission electron microscope (Thermo Fisher Scientific) at a nominal magnification of 36,000 x, corresponding to a super-resolution pixel size of 0.561 Å/px. Automated data collection was performed using SerialEM^79^. 100 frame movies were collected with a Gatan K3 Direct Detection Camera in super-resolution counting mode using a nominal defocus range of 0.8 to 2.0 µm under focus and an estimated dose of 50 e^-^/Å^2^. A total of 15,844 micrographs were collected.

Dose fractioned movies were gain-normalized, motion-corrected, dose-weighted, and binned 2×2 using Fourier cropping using MotionCor2^80^ within the RELION-4.0 wrapper^81^. Motion corrected movies were imported into cryoSPARC v.4.7 and processed using Patch CTF^82^. 2,801,154 particles were initially picked using a three-dimensional IRES-bound-48S reference template^30^, extracted in a 480 px box, and binned to a 128 px box for subsequent processing. Particles were subjected to multiple rounds of reference-free 2D alignment and classification, yielding 741,505 ribosome particles. Iterative 3D focused classification using masks of the ternary complex, eIF1A, and eIF5, resulted in 130,731 particles that contained the initiation factors. These particles were re-extracted without binning in a 480 px box, followed by Bayesian polishing in RELION to correct for beam-induced motion. Polished particles were imported and refined in cryoSPARC. The final particle stack consisting of 129,349 particles yielded a density map of the initiation complex at global resolution of 2.67 Å. Local refinement for the 40S head, body, and ternary complex were conducted separately using the same consensus map to yield final reconstructions maps at 2.58 Å, 2.62 Å, and 2.62 Å, respectively. Masks for 3D focused classification and local refinement were generated using UCSF ChimeraX and cryoSPARC. Figures showing EM maps were generated using UCSF ChimeraX^83^. The processing pipelines are outlined in **Extended Data Fig. 1**.

### Model building, fitting, and refinement

An initial model consisting of the human 48S translation initiation complex from PDB 8OZ0, HCV IRES from PDB 7SYR, and rabbit eIF2 from PDB 6YAL was manually docked in the consensus density map using UCSF ChimeraX. Real space refinement was performed using Phenix^84^. Local refinement and model building were performed using Coot v.1.1^85^ and ISOLDE^86^.

### System setup for molecular dynamics simulations

Molecular dynamics simulations were initiated from the cryo-EM model of the eIF5-bound initiation complex.

To reduce computational cost, a reduced ribosome model in which atoms within 40 Å from the eIF2-tRNA_i_ ternary complex, eIF1A, eIF5, and mRNA molecules are only modeled^87,88^. Heavy atoms 25 Å away from these molecules were harmonically restrained by a force constant of 10 kcal/mol*Å^2^. N- and C-terminal amino acids were capped with acetyl or methyl amide groups, respectively. The reduced ribosome model was solvated in a truncated octahedron of OPC (optimal point charge) water molecules^89^ and 150 mM KCl. Additional K^+^ ions were added to neutralize the system charge. The final system contained ∼361,000 atoms enclosed in a truncated octahedron periodic box defined by lattice vectors lengths of 161.93 Å and interaxial angles of 109.47° (**Extended Data Fig. 5A**). The simulations were parameterized using AMBER ff19SB force field for protein atoms^90^, OL3 with Steinbrecher and Case phosphate oxygens force field for RNA atoms^91^, OPC water model^89^, and 12-6-4-Li Merz ion parameters^92^.

All simulations were performed using OpenMM8.2.0 package^93^ employing periodic boundary conditions, hydrogen mass repartition^94^. Systems were energy minimized for 20,000 steps and then gradually heated to 300 K with harmonic restraints applied to the protein and RNA backbone. Production simulations were performed using a Langevin integrator with a 4 fs timestep and 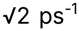 collision rate. Pressure was maintained at 1 atm using a Monte Carlo barostat with an update frequency of 100 steps. Nonbonded interactions were calculated with a 10 Å distance cutoff. For each start site (AUG, CUG, GUG, ACG, UUG, AUC) and eIF5 variant (N30Q, N30K, G29AG31A), eight independent repeats of 2.3 µs in length, each with different initial velocities, were carried out (**Supplementary Table 6**). Trajectory frames were saved every 100 ps. Simulations were performed on the Fred Hutch Scientific Computing cluster using NVIDIA L40S GPUs and mixed numerical precision.

### Trajectory analysis

Trajectory processing was performed with in-house scripts utilizing CPPTRAJ^95^. The last 1.0 µs of each simulation trajectory was considered for analysis. RMSD measurements were performed using the initial structure as reference. Hydrogen bonds were calculated using a 3.2 Å distance cutoff and 120° angle cutoff. Trajectories were visualized using VMD1.9.4^96^.

### Statistical analyses

Unless noted, all analyses were performed using MATLAB (v. R2023-b). E_FRET_ distributions were analyzed as noted above, and we reported the mean ± standard deviation derived from fits of the data to gaussian functions (cftool).

Population-weighted mean binding lifetimes (i.e., the duration of the entire binding event) and FRET state lifetimes (i.e., the duration the complex resided in the low- or high-FRET state during individual events) were calculated using the formula below and reported in the text, where *a* is the fraction of the total population represented by rate *b* (s^-1^), and *c* (s^-1^) is the second rate:

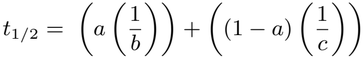

The reported errors were propagated using the formula below, where *a, b*, and *c* are defined as in the equation above, and *e*_*b*_ and *e*_*c*_ are the respective errors of the rates derived from the 95 % C.I. of the fits.

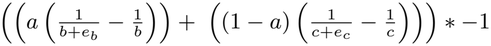

All rates, their respective 95 % C.I., and the number of binding events and complexes analyzed are collated in **Supplementary Table 2**.

The reported errors for mean binding lifetimes of eIF5 bindings events that were on average (mean) low FRET (E_FRET_ ≤ 0.65) or high FRET (E_FRET_ ≥ 0.66) represent 95 % C.I. determined using bootstrapping analyses (n = 2000) and a per molecule resampling strategy, since events on individual complexes were not independent. The number of complexes and binding events analyzed, the calculated means, and 95% C.I. are collated in **Supplementary Table 4**.

For the extract-based translation assays, two-tailed Welch’s t-tests were used to determine statistical significance (n=6, p<0.05).

The reported errors for the association and dissociation conformations represent 95 % C.I. determined using boot-strapping analyses (n = 2000) and a per molecule resampling strategy, since events on individual complexes were not independent.

For molecular dynamics analyses, statistical significance was computed using one-way ANOVA with Dunnett’s multiple comparisons test.

### Use of generative artificial intelligence

We used Anthropic Claude (Opus 5/5.5 and Sonnet 5/5.5) models to help revise the manuscript text and analysis code. All content and data in the manuscript and in the associated GitHub repository were manually reviewed for correctness by the corresponding authors.

---

**Extended Data Figure 1:**
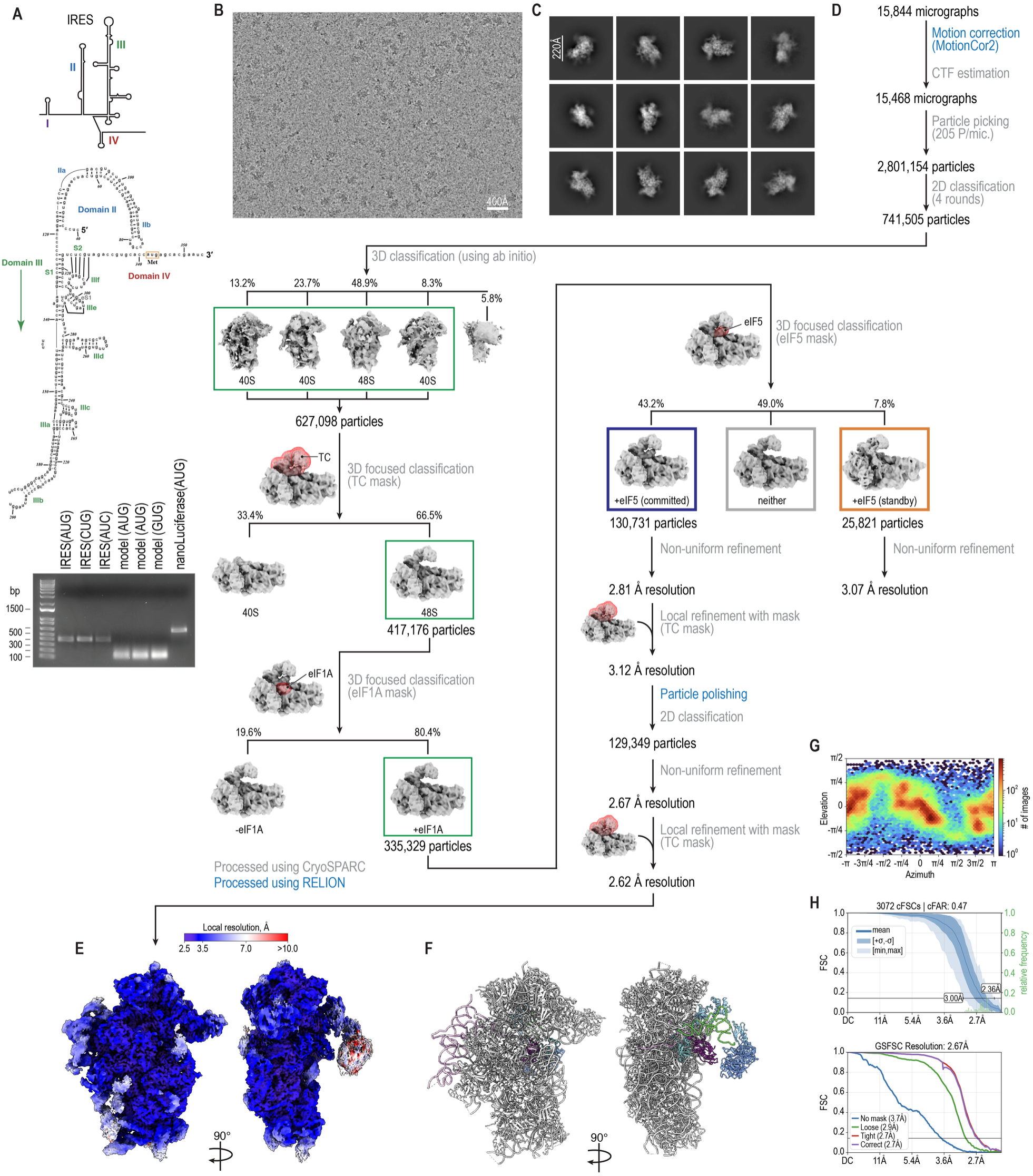
Cryo-EM processing of minimal initiation complex. **A**, Diagram of the hepatitis C virus internal ribosome entry site (IRES) structure (top, middle) and a representative gel image after analysis of representative IRES, unstructured model, and nanoluciferase mRNAs used in this study (bottom). **B**, Representative micrograph of minimal initiation complex bound to the IRES with an AUG start site. **C**, Representative 2D class averages of initiation complexes. **D**, Processing pipeline highlighting the major classification and refinement steps. Particle counts and class population for each significant step are shown. **E**, Local resolution estimation of the minimal initiation complex. **F**, Atomic model of the minimal initiation built from the density map. **G**, Angular distribution of particles used in the final reconstruction. **H**, Three-dimensional conical FSC curves (top) and gold-standard FSC curves (bottom) used in the final reconstruction.

**Extended Data Figure 2:**
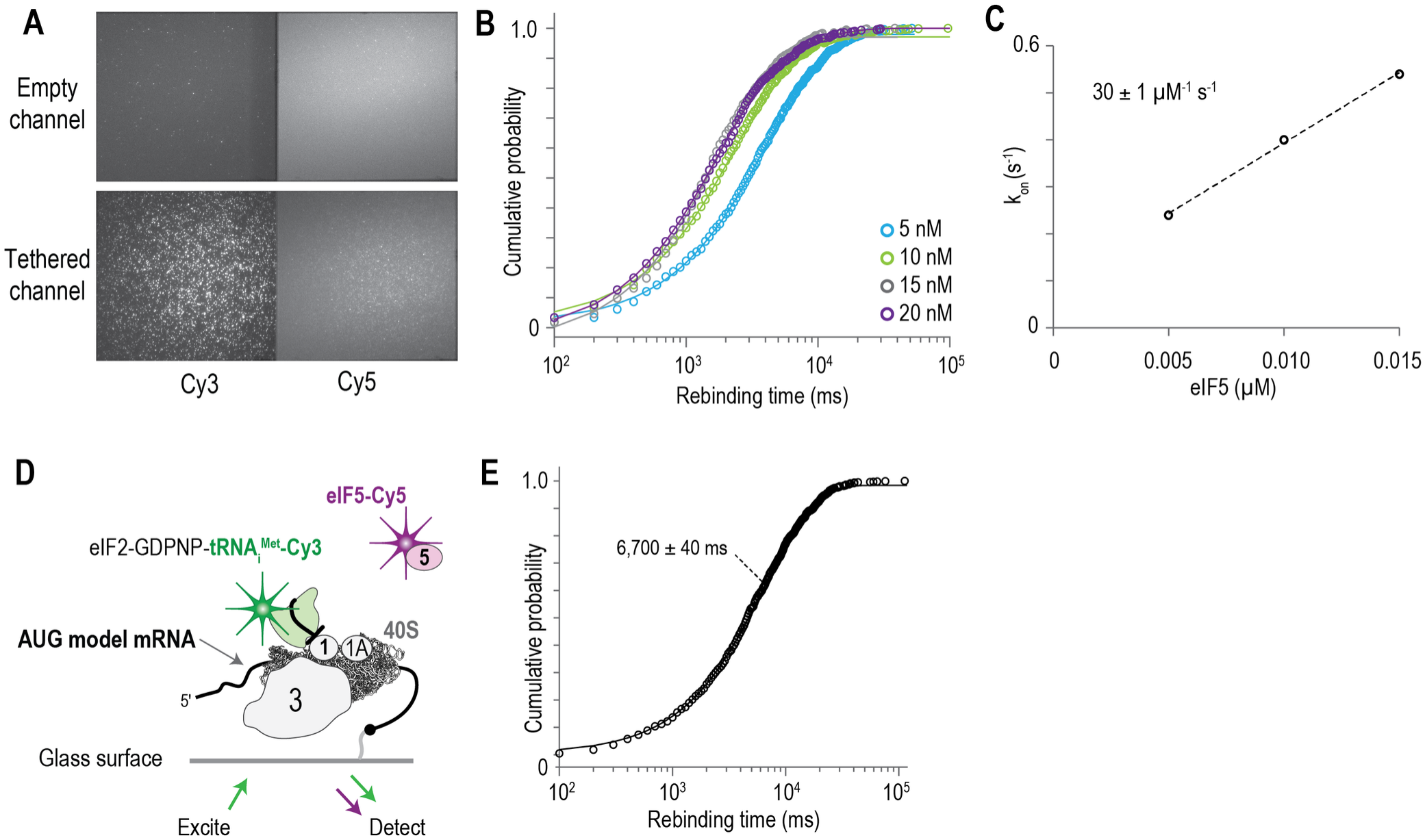
Direct analysis of eIF5 binding at start codons. **A**, Example fields of view of TIRFm experiments. **B**, Cumulative probability plot of eIF5 rebinding times at the indicated eIF5 concentration. Lines represent fits to exponential functions. **C**, Plot of the observed eIF5 association rate constants at the indicated concentrations. The line represents a fit by linear regression analysis to derive the indicated association rate. **D**, Schematic of the single-molecule human translation initiation assay using the model mRNA and all eIFs (except eIF4 proteins). **E**, Cumulative probability plot of the rebinding time. Line represents fit to a single-exponential function and the derived mean time is reported on the plot.

**Extended Data Figure 3:**
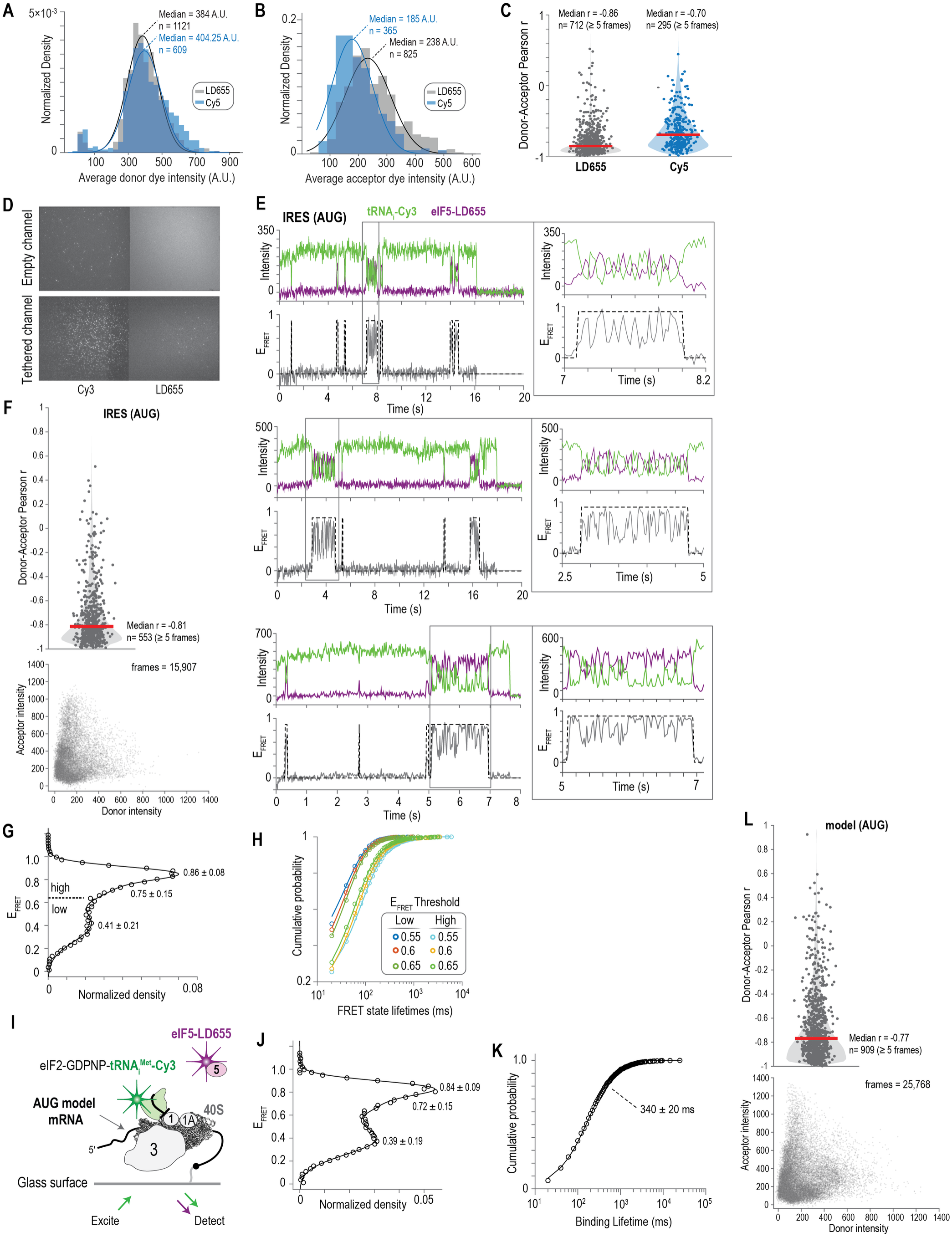
Rapid and reversible sampling of multiple conformations. **A**, Histogram of the Cy3 donor dye intensities observed outside of eIF5 binding events (FRET off state) in experiments with Cy5-eIF5 and LD655-eIF5 present. The line represents a fit to a gaussian function, with the medians and numbers of analyzed events indicated. **B**, Histogram of either Cy5 or LD655 acceptor dye intensities observed during eIF5 binding events. The line represents a fit to a gaussian function, with the medians and numbers of analyzed events indicated. **C**, Violin plot of the calculated correlation between donor and acceptor fluorescence intensities during individual eIF5 binding events using the indicated acceptor dyes (LD655 or Cy5), where r is the Pearson’s correlation coefficient and n is the number of binding events that last 5 frames or longer. The red line indicates the median r. **D**, Example fields of view of TIRFm experiments at 20 ms temporal resolution. **E**, Example single-molecule fluorescence data collected at 20 ms temporal resolution, where tRNAi–Cy3 (green) and eIF5–LD655 (purple) were monitored during translation initiation on the IRES with an AUG start site. The dashed line demarcates eIF5 binding events. Zoomed in portions of the fluorescence data show rapid fluctuations between FRET states within individual binding events. **F**, Violin plot of the calculated correlation between donor and acceptor fluorescence intensities during individual eIF5 binding events, where r is the Pearson’s correlation coefficient and n is the number of binding events that last 5 frames or longer (top). Scatter plot of the acceptor and donor intensities measured in each frame (20 ms exposures) during eIF5 binding events (bottom). **G**, tRNA_i_-Cy3 – to – eIF5-LD655 FRET efficiency distribution; the line represents a fit to a double gaussian function with the indicated means ± standard deviation. The dashed line indicates the defined E_FRET_ threshold that separates the low- and high-FRET states. **H**, Cumulative probability plot of the FRET state lifetimes as determined using the indicated E_FRET_ threshold. Lines represent fits to double exponential functions. **I**, Schematic of the single-molecule human translation initiation assay using the model human mRNA and all eIFs (except eIF4 proteins). **J**, tRNA_i_-Cy3 – to – eIF5-LD655 FRET efficiency distribution using the model human mRNA; the line represents a fit to a double gaussian function with the indicated means ± standard deviation. **K**, Cumulative probability plot of the eIF5 binding lifetime. Line represents fit to a double-exponential function and the derived mean time is reported on the plot. **L**, Violin plot of the calculated correlation between the donor and acceptor fluorescence intensities during individual eIF5 binding events, where r is the Pearson’s correlation coefficient and n is the number of binding events that last 5 frames or longer (top). Scatter plot of the acceptor and donor intensities measured in each frame (20 ms exposures) during eIF5 binding events (bottom).

**Extended Data Figure 4:**
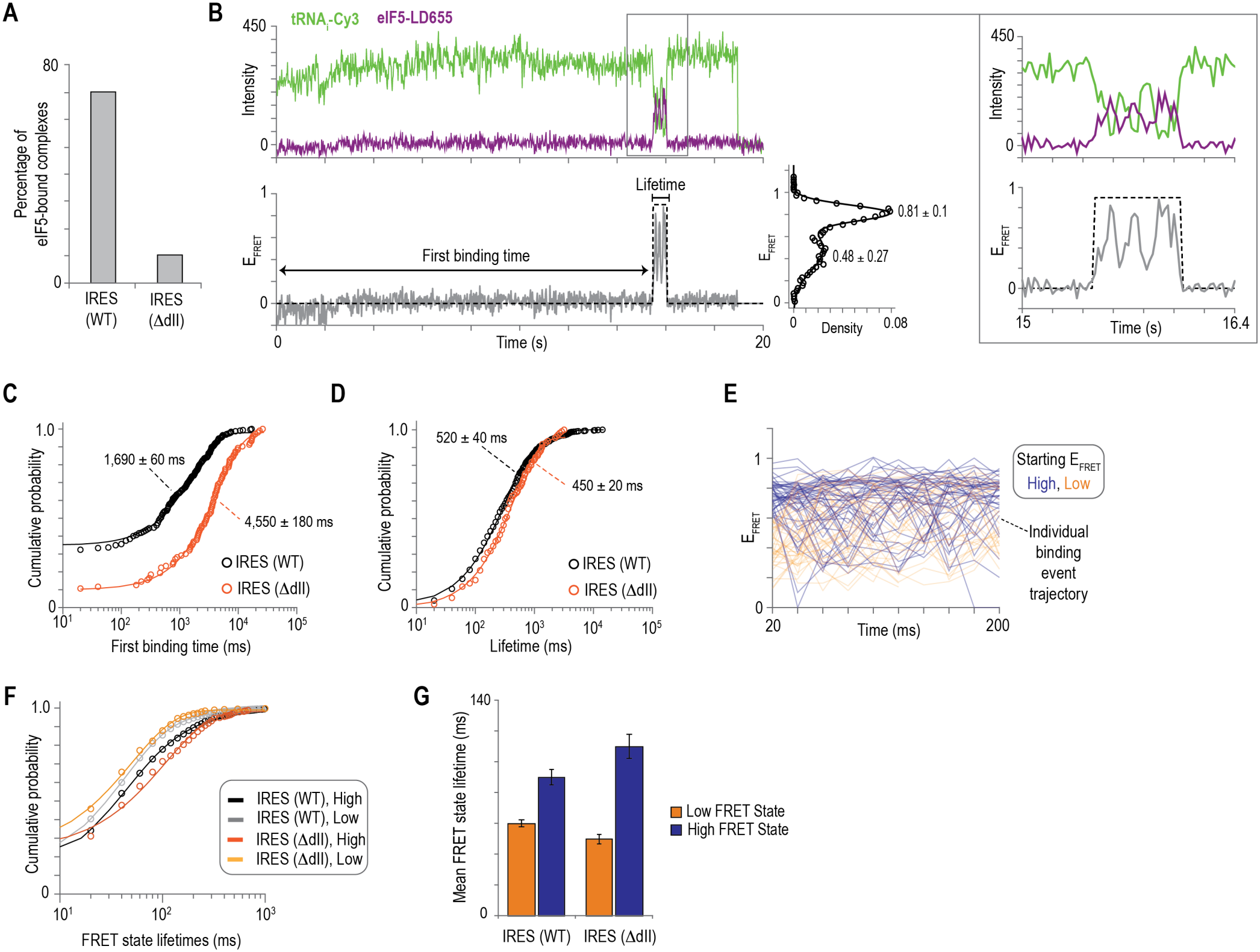
Rapid fluctuations occur independent of domain II of the IRES. **A**, Percentage of eIF5-bound complexes observed on the wild-type IRES and the IRES with the domain II deletion (ΔdII). **B**, Example single-molecule fluorescence data collected at 20 ms temporal resolution, where tRNAi–Cy3 (green) and eIF5–LD655 (purple) were monitored during translation initiation on the IRES (ΔdII). The dashed line demarcates eIF5 binding events. The corresponding FRET efficiency distribution reports the FRET values of each population; the line represents a fit to a double gaussian function with the indicated means ± standard deviation. The zoomed in portion of the fluorescence data shows rapid fluctuations between FRET states within an individual binding event. **C**, Cumulative probability plot of the time between starting data collection and the first eIF5 binding event observed with the wild-type IRES and the IRES (ΔdII). Lines represent fits to single-exponential functions and the derived mean times are reported on the plot. **D**, Cumulative probability plot of eIF5 binding lifetimes during translation initiation on the wild type and (ΔdII) IRES. Lines represent fits to exponential functions and the derived mean times are reported on the plot. **E**, Plot of the observed FRET efficiency trajectories of two hundred binding events over the first two hundred milliseconds. Blue lines represent events that started in high FRET (≥0.66) and orange lines represent events that started in low FRET (≤ 0.65). **F**, Cumulative probability plot of the high- and low-FRET state lifetimes for the wild-type and (ΔdII) IRES. Lines represent fits to exponential functions. **G**, Weighted population means of the high- and low-FRET state lifetimes during translation initiation on the wild-type and (ΔdII) IRES.

**Extended Data Figure 5:**
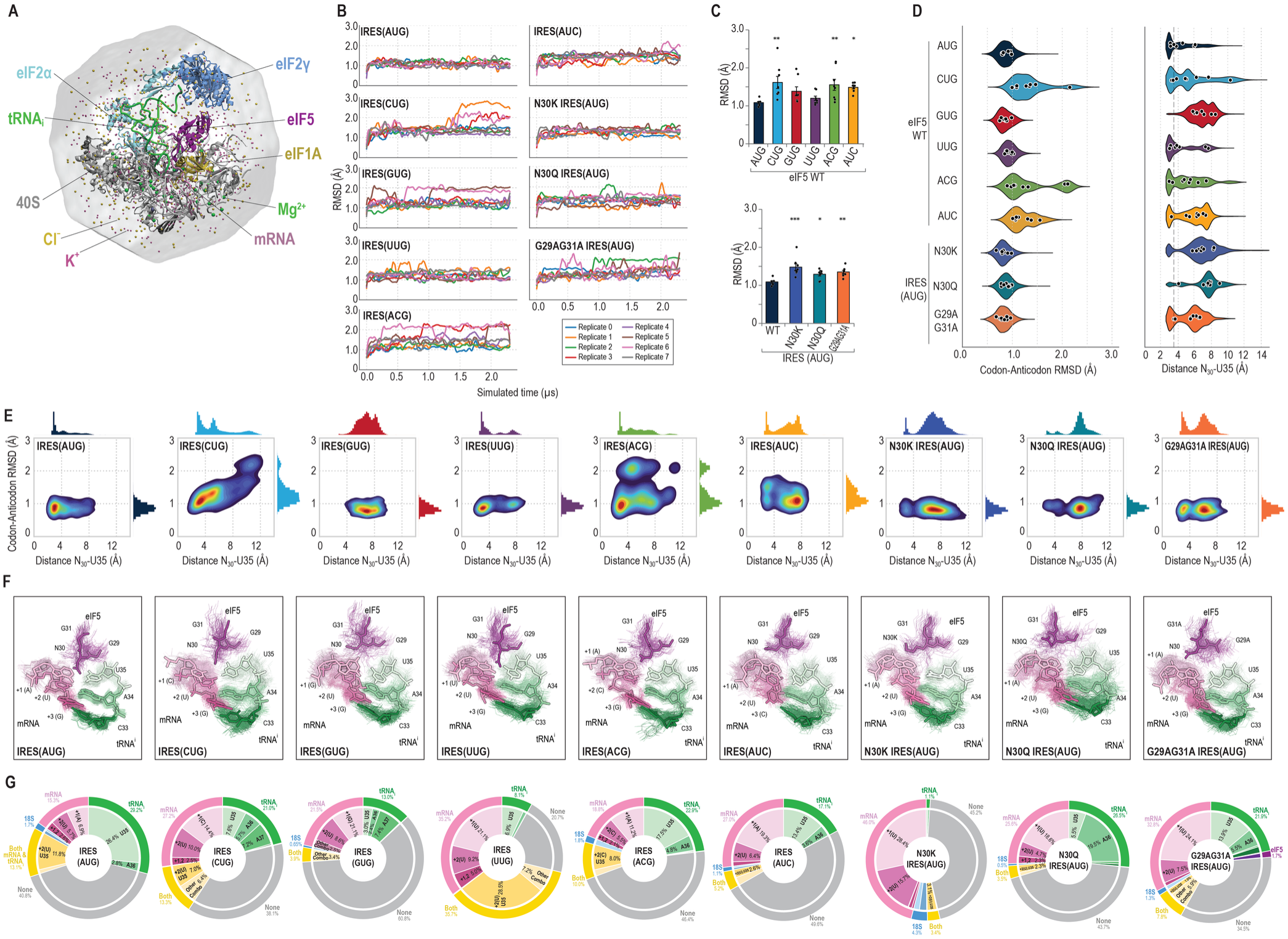
Molecular dynamics simulation of reduced ribosome model reveals rearrangement of hydrogen bonds. **A**, Initial molecular dynamics snapshot of the reduced ribosome model solvated in a truncated octahedron. **B**, Time-resolved RMSD of the hydrogen bond network involving the translation start site nucleotides (mRNA), anticodon nucleotides (tRNA), and G_29_N_30_G_31_ loop (eIF5) at different translation start sites. RMSD is calculated using the initial structure as a reference. Individual replicates are colored accordingly. **C**, Average RMSD of the hydrogen bond network from (**B**). Values are calculated from the last 1.0 µs of simulations across 8 replicates. Error bars represent SEM. Statistical significance is indicated by * p < 0.05, and ** p <0.01 (one-way ANOVA with Dunnett’s multiple comparisons test versus the wild type). **D**, Codon-anticodon heavy atom RMSD and N30-U35 distance distribution of the trajectory frames from the last 1.0 µs of simulations. Median value of each replicate is shown as black circles. **E**, Conformational landscape of N_30_-U35 distance and codon-anticodon RMSD distributions from (D). **F**, Molecular dynamics snapshots of the codon-anticodon interface. Snapshots were obtained at 100-ns intervals from the last 1.0 µs of each of eight simulation replicates (80 snapshots). The representative structure, shown as opaque, is from the most populated cluster of the conformational landscape in (**E**). **G**, Hydrogen bond occupancy of the N_30_ side chain amide calculated as the percentage of simulation frames assigned to mutually exclusive interaction states for each translation state site. Individual residues corresponding to each molecule are indicated.

**Extended Data Figure 6:**
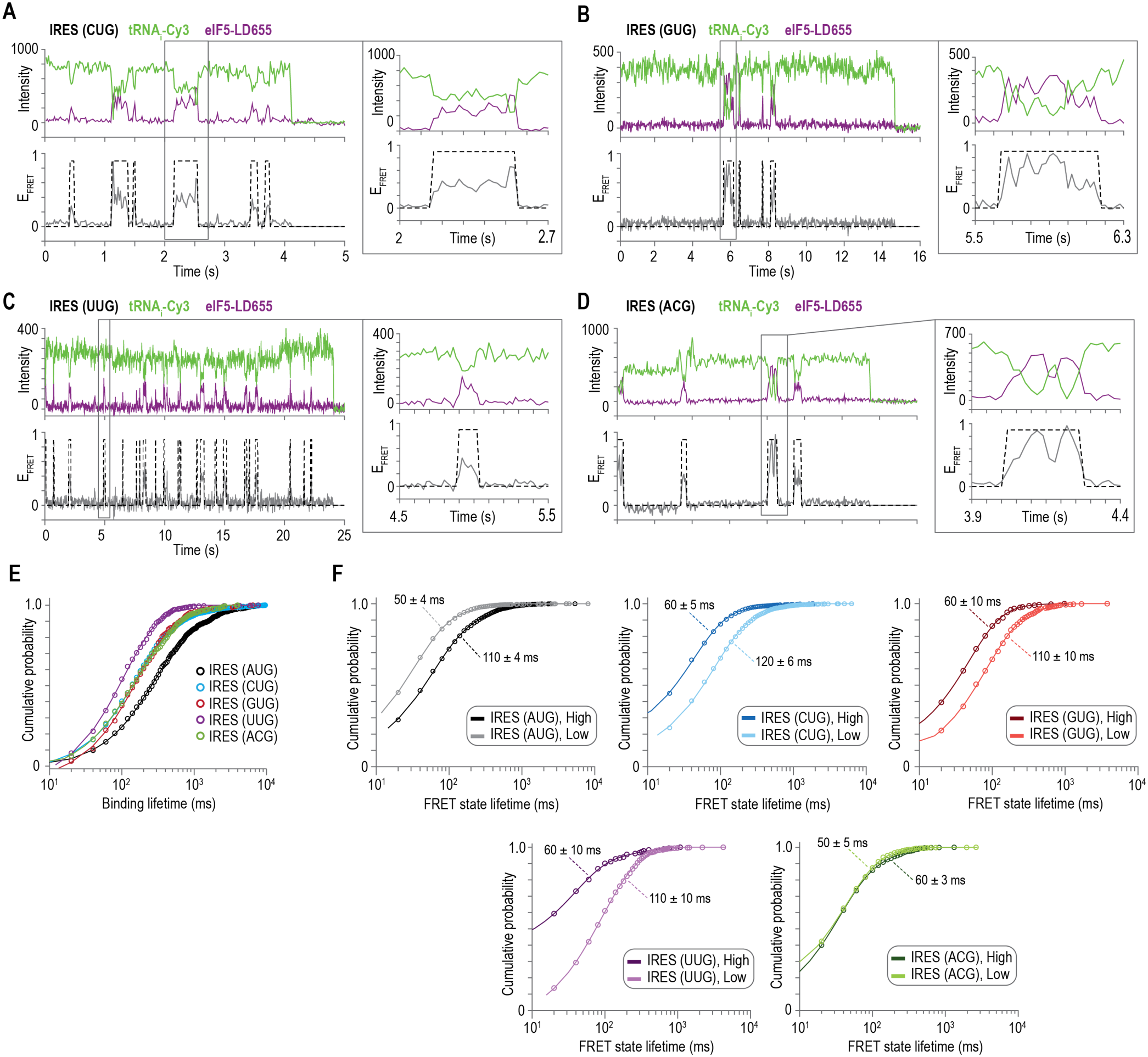
Start codon identity biases conformational occupancy. **A-D**, Example single-molecule fluorescence data collected at 20 ms temporal resolution, where tRNAi–Cy3 (green) and eIF5–LD655 (purple) were monitored during translation initiation on the indicated IRES start site variant. The dashed line demarcates eIF5 binding events. Zoomed in portions of the fluorescence data show varying amounts of rapid fluctuations between FRET states within individual binding events. **E**, Cumulative probability plot of eIF5 binding lifetimes during translation initiation on the indicated IRES variants. Lines represent fits to double-exponential functions. **F**, Cumulative probability plots of the high- and low-FRET state lifetimes during eIF5 binding at each IRES variant. Lines represent fits to exponential functions and the derived mean times ± 95 % CI are reported on the plots.

**Extended Data Figure 7:**
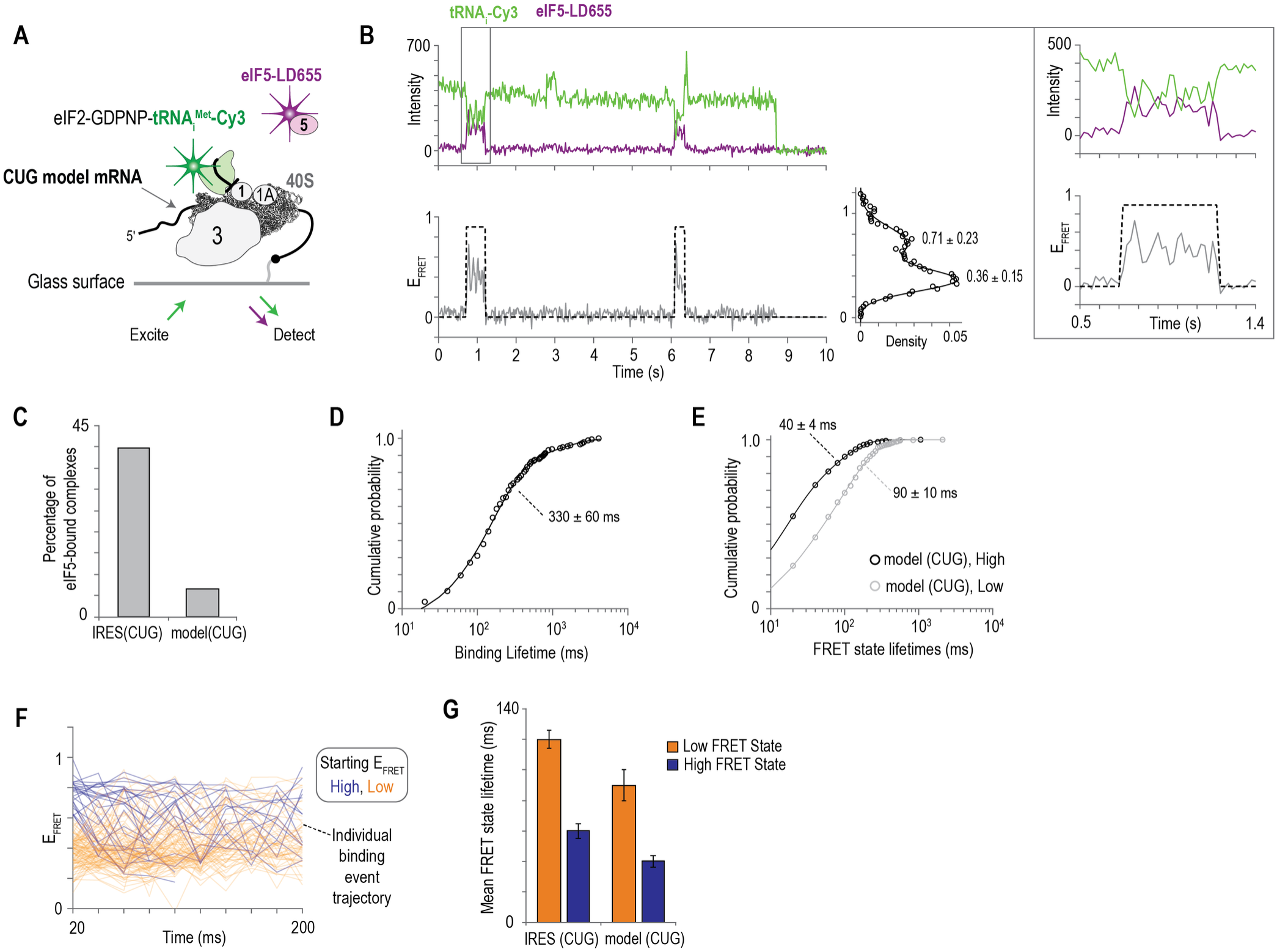
eIF5 binding to canonical initiation complexes formed on an unstructured model mRNA with a CUG start site. **A**, Schematic of the single-molecule human translation initiation assay using the model mRNA with a CUG start site and all eIFs (except eIF4 proteins). **B**, Example single-molecule fluorescence data collected at 20 ms temporal resolution, where tRNAi–Cy3 (green) and eIF5–LD655 (purple) were monitored during translation initiation on the model CUG mRNA. The dashed line demarcates eIF5 binding events. The corresponding FRET efficiency distribution reports the FRET values of each population; the line represents a fit to a double gaussian function with the indicated means ± standard deviation. Zoomed in portion of the fluorescence data shows reduction of rapid fluctuations between FRET states within an individual binding event. **C**, Percentage of eIF5-bound complexes observed on the IRES (CUG) and the CUG model mRNA. **D**, Cumulative probability plot of the eIF5 binding lifetime. Line represents fit to a double-exponential function and the derived mean time is reported on the plot. **E**, Cumulative probability plot of the high- and low-FRET state lifetimes. Lines represent fits to exponential functions and the derived mean times ± 95 % CI are reported on the plots. **F**, Plot of the observed FRET efficiency trajectories of two hundred binding events over the first two hundred milliseconds. Blue lines represent events that started in high FRET (≥0.66) and orange lines represent events that started in low FRET (≤ 0.65). **G**, Weighted population means of the high- and low-FRET state lifetimes during translation initiation on the IRES (CUG) and the CUG model mRNA.

**Extended Data Figure 8:**
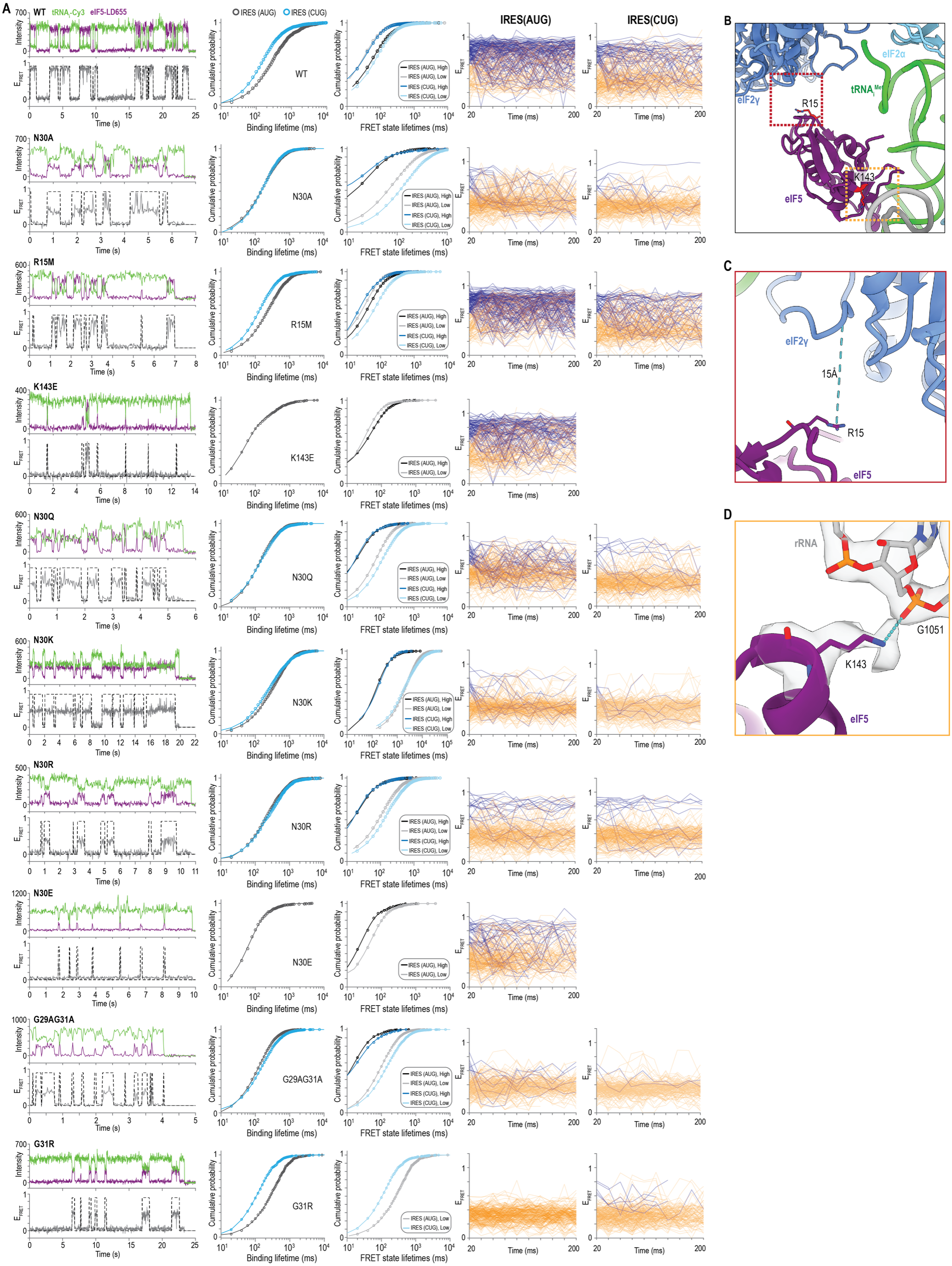
A conserved eIF5 loop controls codon sensitivity. **A**, For each eIF5 variant, columns from left to right show: Example single-molecule fluorescence data collected at 20 ms temporal resolution, where tRNAi–Cy3 (green) and eIF5–LD655 (purple) were monitored during translation initiation on IRES (AUG). The dashed line demarcates eIF5 binding events. Cumulative probability plot of eIF5 binding lifetimes during translation initiation on IRES (AUG) and IRES (CUG). Lines represent fits to double-exponential functions. Cumulative probability plots of the high- and low-FRET state lifetimes during translation initiation on IRES (AUG) and IRES (CUG). Lines represent fits to double-exponential functions. Plots of the observed FRET efficiency trajectories of two hundred binding events over the first two hundred milliseconds. Blue lines represent events that started in high FRET (≥0.66) and orange lines represent events that started in low FRET (≤ 0.65). **B-D**, Atomic model of minimal initiation complex highlighting sites of eIF5 substitutions. **B**,**C** Views of eIF5 R_15_ in proximity to eIF2γ. Distance between eIF5 R_15_ to eIF2γ I_39_ used as a proxy for protein–protein proximity. **D**, Close-up view of eIF5 K_143_ interacting with rRNA. The grey surface represents the electron density map, with local resolution of 3.2 Å.

**Extended Data Figure 9:**
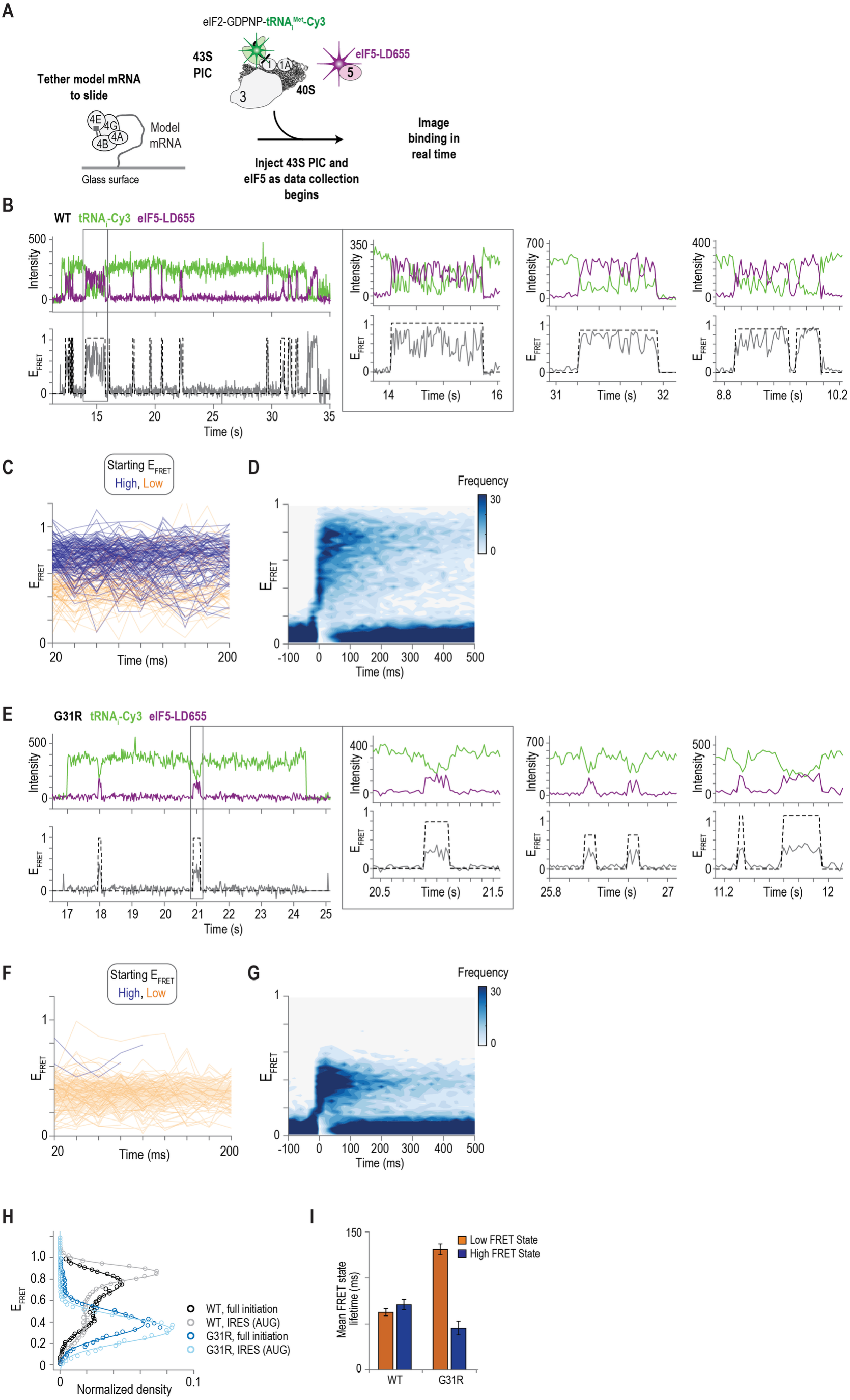
eIF5 rapidly and reversibly fluctuates between low- and high-FRET states after 43S loading and scanning in real time. **A**, Schematic of the real-time full initiation single-molecule assay using eIF2-GDPNP and stopped-flow injection of eIF5-LD655. **B**,**E** Example single-molecule fluorescence data collected at 20 ms temporal resolution, where tRNAi–Cy3 (green) and eIF5– LD655 (purple) were monitored during translation initiation on the model mRNA with an AUG start site. The dashed line demarcates eIF5 binding events. The corresponding FRET efficiency distribution reports the FRET values of each population. Zoomed in portion of the fluorescence data shows rapid fluctuations between FRET states within an individual binding event. **C**,**F** Plots of the observed FRET efficiency trajectories of two hundred binding events over the first two hundred milliseconds. Blue lines represent events that started in high FRET (≥0.66) and orange lines represent events that started in low FRET (≤ 0.65). **D**,**G** Density plot of the observed FRET Efficiency (E_FRET_) synchronized to the start of all wild-type eIF5 binding events (831) observed on 408 complexes and all eIF5^G31R^ binding events (846) observed on 207 complexes. **H**, tRNA_i_-Cy3 – to – eIF5-LD655 FRET efficiency distributions observed for wild-type eIF5 and eIF5^G31R^; the lines represent fits to double gaussian functions. **I**, Weighted population means of the high- and low-FRET state lifetimes during translation initiation of wild-type eIF5 and eIF5^G31R^. Panels B-D correspond to wild-type eIF5. Panels E-G correspond to the G31R variant.

**Extended Data Figure 10:**
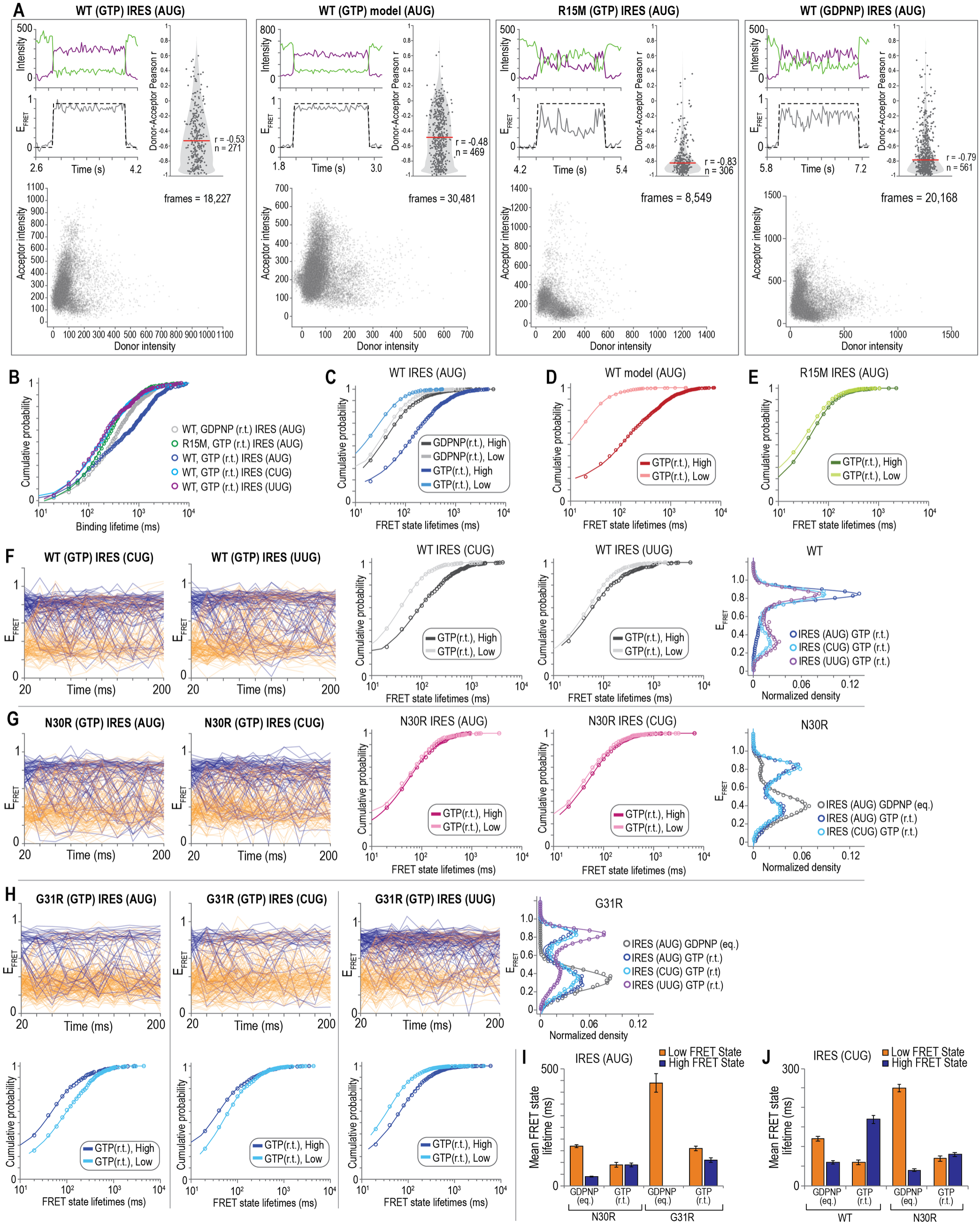
GTP hydrolysis stabilizes the high-FRET conformation. **A**, For each indicated condition: Example single-molecule fluorescence data collected at 20 ms temporal resolution where the dashed line demarcates eIF5 binding events. Violin plot of the calculated correlation between donor and acceptor fluorescence intensities during individual eIF5 binding events, where r is the Pearson’s correlation coefficient and n is the number of binding events that last 5 frames or longer. Scatter plot of the acceptor and donor intensities measured in each frame (20 ms exposures) during eIF5 binding events. **B**, Cumulative probability plot of the binding lifetimes for each of the conditions examined. Lines represent fits to double-exponential functions. **C**, Cumulative probability plot of the high- and low-FRET state lifetimes during wild-type eIF5 binding in the presence of GDPNP and GTP. Lines represent fits to double-exponential functions. **D**, Cumulative probability plot of the high- and low-FRET state lifetimes during wild-type eIF5 binding canonical initiation complexes assembled on a model RNA in the presence of GTP. Lines represent fits to double-exponential functions. **E**, Cumulative probability plot of the high- and low-FRET state lifetimes during eIF5^R15M^ binding in the presence of GTP. Lines represent fits to double-exponential functions. **F-G**, For wild-type (**F**) and eIF5^N30R^ (**G**) from left to right: For each IRES start site variant indicated, plots of the observed FRET efficiency trajectories of two hundred binding events over the first two hundred milliseconds. Blue lines represent events that started in high FRET (≥0.66) and orange lines represent events that started in low FRET (≤ 0.65). Cumulative probability plots of the high- and low-FRET state lifetimes. Lines represent fits to double-exponential functions. tRNA_i_-Cy3 – to – eIF5-LD655 FRET efficiency distributions for each of the indicated conditions; the lines represent fits to double gaussian functions. **H**, For eIF5^G31R^ binding at each indicated IRES start site variant: plots of the observed FRET efficiency trajectories of two hundred binding events over the first two hundred milliseconds. Blue lines represent events that started in high FRET (≥0.66) and orange lines represent events that started in low FRET (≤ 0.65). Cumulative probability plots of the high- and low-FRET state lifetimes. Lines represent fits to double-exponential functions (bottom). tRNA_i_-Cy3 – to – eIF5-LD655 FRET efficiency population distributions for each of the indicated conditions (right). The corresponding FRET efficiency distribution reports the FRET values of each population; the lines represent fits to double gaussian functions. **I**, Weighted population means of the high- and low-FRET state lifetimes for eIF5^N30R^ and eIF5^G31R^ at the AUG start site in the presence of GDPNP or GTP. **J**, Weighted population means of the high- and low-FRET state lifetimes for wild-type eIF5 and eIF5^N30R^ at the CUG start site in the presence of GDPNP or GTP.

**Extended Data Figure 11:**
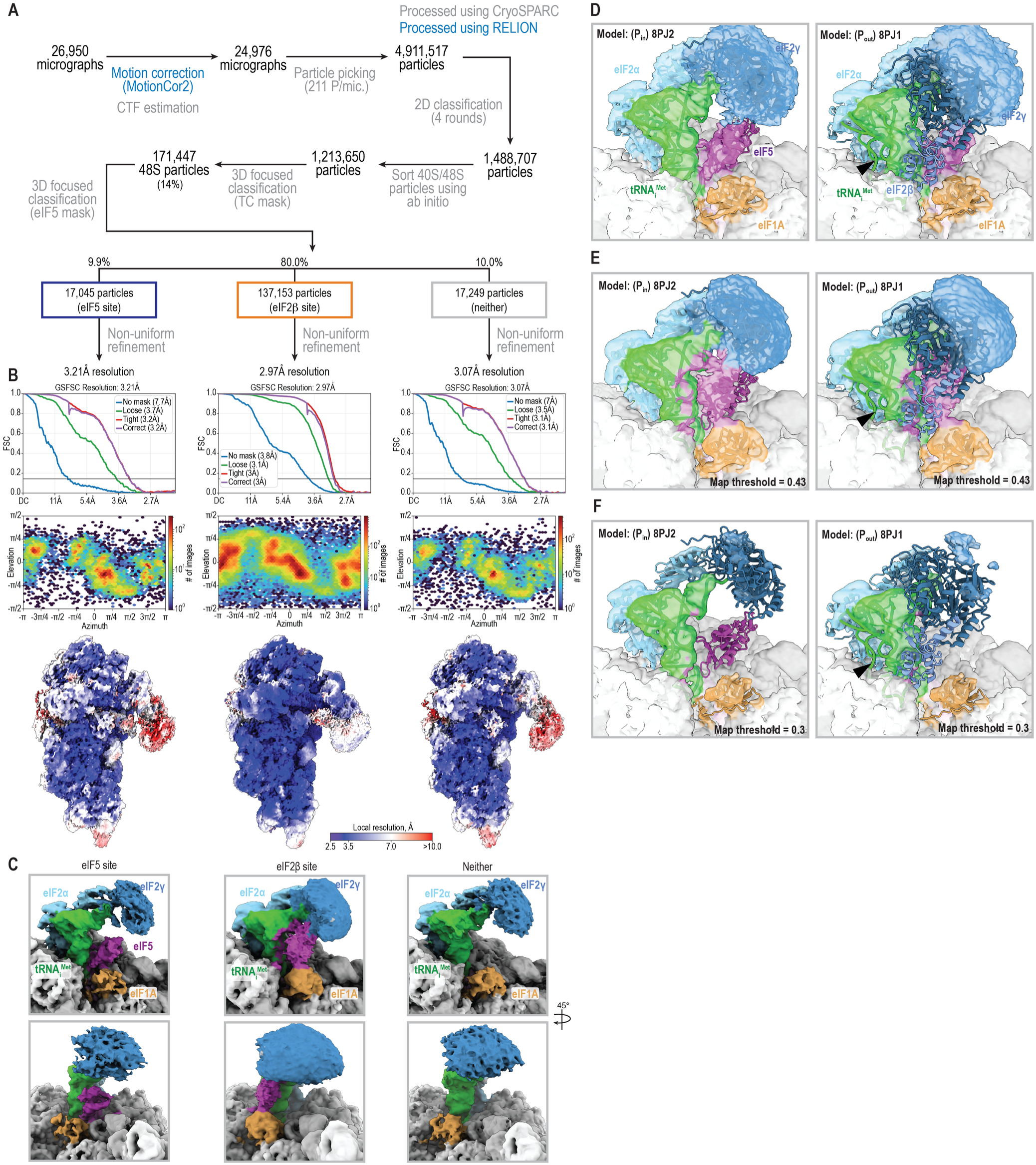
Cryo-EM analysis of the minimal initiation complex on UUG start codon. **A**, Processing pipeline highlighting the major classification and refinement steps for the initiation complex on UUG start codon. Particle counts and class population for each significant step are shown. **B**, Three-dimensional gold standard FSC curves, angular distribution of particles, and local resolution estimation of the three resolved classes. **C**, Cryo-EM density maps of the three classes of 48S particles, defined as in Fig. 6A. **D-F**, In the UUG dataset, the density of tRNA_i_ from both the eIF5-site and eIF2β-site classes closely resembles the P_in_ state. Atomic models of the initiation complex in the P_in_ state (PDB: 8PJ2) and the P_out_ state (PDB:8PJ1) fitted in the density maps of the eIF5-site class (**D**) and the eIF2β-site class (**E, F**). Density of the eIF2β-site class is shown at lower threshold in panel F.

**Extended Data Figure 12:**
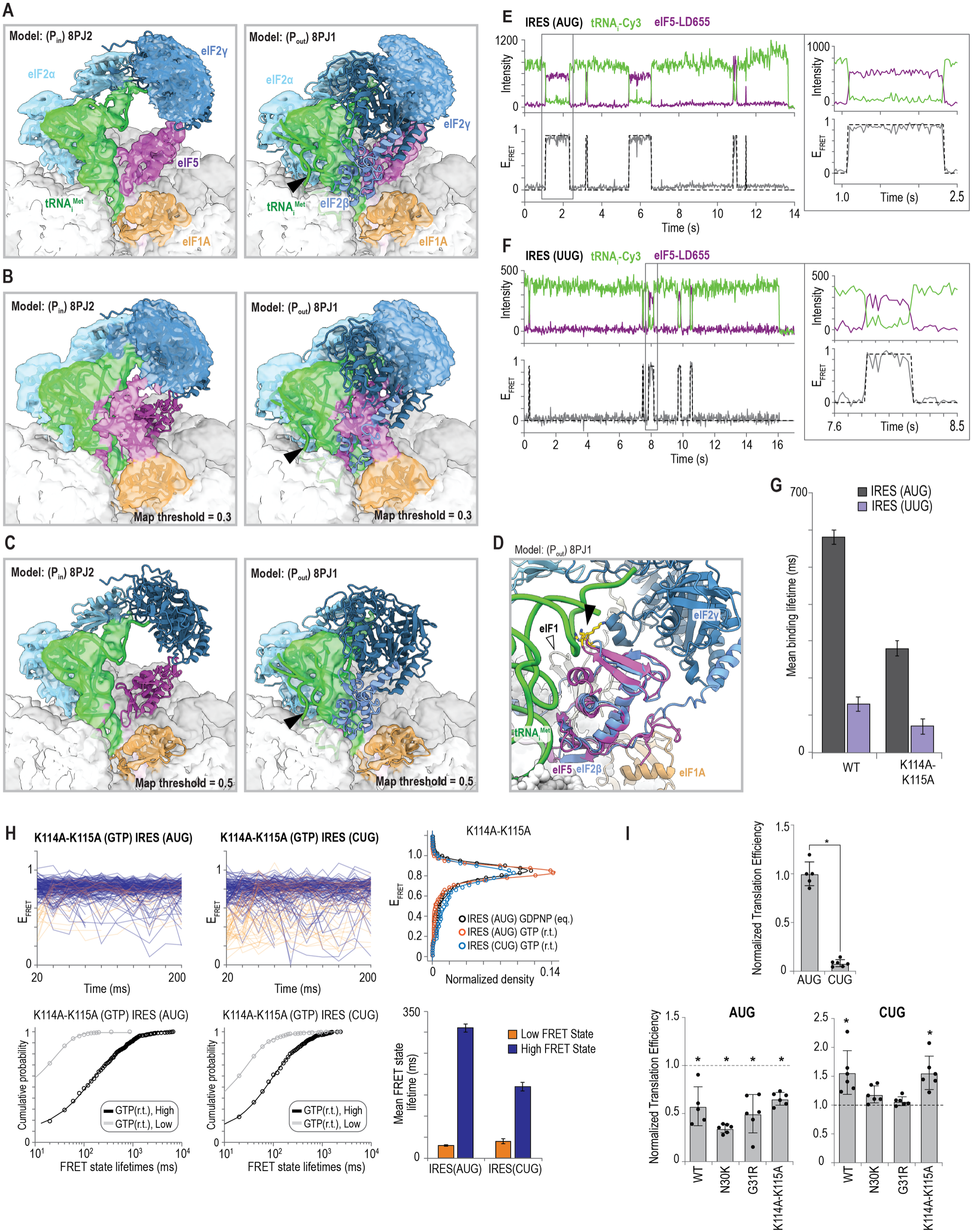
Destabilization of the low-FRET conformation. **A-C**, In the AUG dataset, density of the tRNA_i_ from the eIF5-site and eIF2β-site classes closely resembles the P_in_ state. Atomic models of the initiation complex in the P_in_ state (PDB: 8PJ2) and the P_out_ state (PDB:8PJ1) fitted in the density maps of the eIF5-site class (**A**) and the eIF2β-site class (**B, C**). Density of the eIF2β-site class is shown at lower threshold in panel C. **D**, Atomic model of the P_out_ state (PDB:8PJ1) with eIF5 N-terminal domain (purple) superimposed on eIF2β core domain (cornflower blue). Residues R_296_ of eIF2β and analogous residues K_114_ and K_115_ of eIF5 are indicated with the black triangle eIF1 is indicated with the white triangle. **E-F**, Example single-molecule fluorescence data collected at 20 ms temporal resolution, where tRNAi–Cy3 (green) and eIF5^K114A-K115A^–LD655 (purple) were monitored during translation initiation on the indicated IRES start site variant. The dashed line demarcates eIF5 binding events. Zoomed in portions of the fluorescence data show varying amounts of rapid fluctuations between FRET states within individual binding events. **G**, Weighted population means of binding lifetimes for the indicated eIF5 variants. **H**, For each IRES start codon variant indicated, plots of the observed FRET efficiency trajectories of two hundred binding events over the first two hundred milliseconds. Blue lines represent events that started in high FRET (≥ 0.66) and orange lines represent events that started in low FRET (≤ 0.65). Cumulative probability plots of the high- and low-FRET state lifetimes. Lines represent fits to double-exponential functions. tRNA_i_-Cy3 – to – eIF5-LD655 FRET efficiency distributions for each of the indicated conditions; the lines represent fits to double gaussian functions. Weighted population means of the high- and low-FRET state lifetimes for eIF5^K114A-K115A^ at the AUG and CUG start site in the presence of GTP. **I (top)**, Translation efficiency plot of the control reactions for each nLuc reporter start site variant relative to the AUG codon (n=6, p < 0.0001). **I (lower left)**, Translation efficiency plot of the AUG start codon nLuc reporter upon addition of the indicated eIF5 variants relative to the translation efficiency of the control reaction for the AUG nLuc reporter (dashed line) (n=6, p-values from left to right: 0.0057, <0.0001, 0.0008, 0.0013). **I (lower right)**, Translation efficiency plot of the CUG start codon nLuc reporter upon addition of the indicated eIF5 variants relative to the translation efficiency of the control reaction for the CUG nLuc reporter (dashed line) (n=6, p-values from left to right: 0.0431, 0.3834, 0.7507, 0.0344). Asterisks indicate significance (*p*<0.05) determined by a two-tailed Welch’s t-test. Control reactions received an equal volume of eIF5 storage buffer without protein.

**Extended Data Figure 13:**
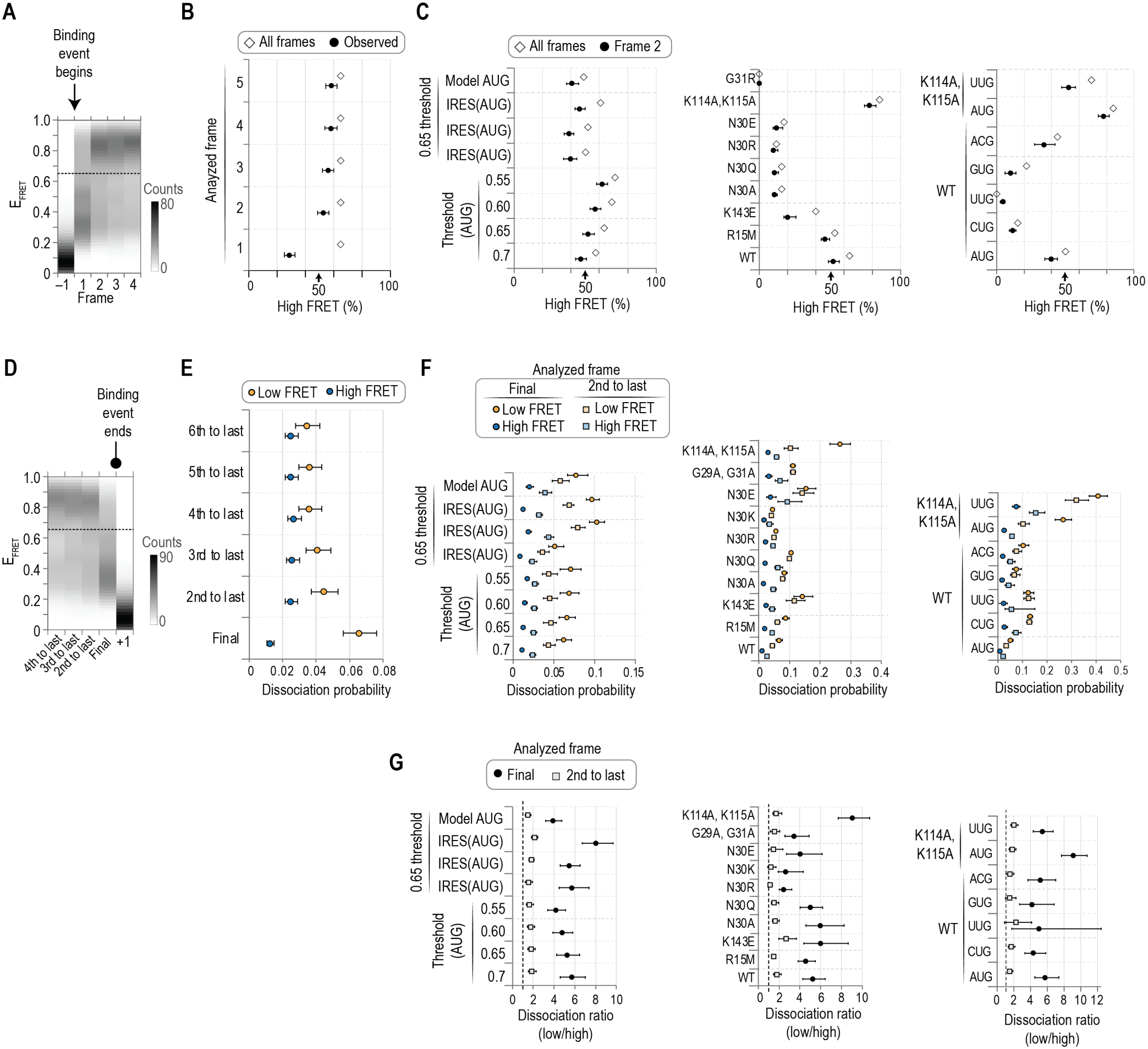
Association and dissociation conformation analyses. **A**,**D** Example density plots of the observed E_FRET_ in the indicated frame. Frame 1 was defined as the frame when wild-type eIF5 bound the initiation complex (i.e., FRET appeared) (panel A), and the ‘Final’ frame as the frame when eIF5 dissociated (FRET abolished) (panel D). Both plots represent wild-type eIF5 binding to initiation complexes poised at an AUG start codon. The dashed lines indicate the defined threshold (0.65) between the low- and high-FRET states. **B**, Plot of the percent of frames that yielded high FRET (≥ 0.66) in the indicated frame (black circles) relative to the percent of all observed frames across all binding events (open diamonds). **C**, Plots of the percent of frames that yielded high FRET as defined using different thresholds and across replicates (left), across the indicated eIF5 variants on an AUG codon (middle), and across the indicated start codons and variants (right). Since frame 1 includes time censored binding events, we focused analyses on frame 2 (see, Methods). **E**, Plot of the eIF5 dissociation probability occurring in either the low-FRET (orange) or high-FRET (blue) states as calculated using the indicated frame as the reference. **F**, Plots of the eIF5 dissociation probability occurring in either the low-FRET (orange) or high-FRET (blue) states as calculated using the final (circles) or second-to-last (squares) as the reference frame. **G**, Plots of the calculated dissociation ratios (low / high). On all plots, the black bars represent the 95% C.I. yielded from bootstrapping analyses of the observed values (n = 2000).

**Supplementary Table 1:** Cryo-EM data collection, refinement, and validation statistics.

| Dataset | **Minimal initiation complex on IRES w/ AUG** |
|---|---|
|  | (EMDB-75872)<br>(PDB 11NV) |
| **Data collection and processing** |  |
| Magnification | 36000x |
| Voltage (kV) | 200 |
| Electron exposure (e–/Å<sup>2</sup>) | 50 |
| Defocus range (µm) | 0.8 – 2.0 under focus |
| Pixel size (Å) | 1.122 |
| Symmetry imposed | C1 |
| Initial particle images (no.) | 2,801,154 |
| Final particle images (no.) | 129,349 |
| Map resolution (Å) | 2.7 |
| FSC threshold | 0.143 |
| Map resolution range (Å) | 2.5 – 41.7 |
| **Refinement** |  |
| Initial model used (PDB code) | 8OZ0 |
| Model resolution (Å) | 3.0 |
| FSC threshold | 0.5 |
| Model resolution range (Å) | 2.5 – 41.7 |
| Map sharpening *B* factor (Å<sup>2</sup>) | -62.3 |
| **Model composition** |  |
| Non-hydrogen atoms | 85442 |
| Protein residues | 5713 |
| Nucleotides | 1861 |
| Ligands | 86 |
| ***B* factors (Å<sup>2</sup>)** |  |
| Protein | 47.86 |
| Nucleotide | 45.37 |
| Ligand | 37.59 |
| **R.m.s. deviations** |  |
| Bond lengths (Å) | 0.002 (1) |
| Bond angles (°) | 0.603 (13) |
| **Validation** |  |
| MolProbity score | 1.43 |
| Clashscore | 4.98 |
| Poor rotamers (%) | 0.83 |
| **Ramachandran plot** |  |
| Favored (%) | 97.07 |
| Allowed (%) | 2.93 |
| Disallowed (%) | 0.00 |
| **RNA** |  |
| Correct sugar puckers (%) | 100.0 |

**Supplementary Table 2: Single-molecule experiments, binding kinetics, and FRET state lifetimes.** Provided as a separate file (`Supplementary Table 2.xlsx`). All rates, their respective 95 % C.I., and the number of binding events and complexes analyzed are collated in this table.

**Supplementary Table 3:** Hydrogen bond lifetime of eIF5 N30 and tRNA_i_ U35.

| eIF5 | Start Site | n trajectories with ≥1 H-bonds observed | Total H-bond events in all trajectories | Lifetime (ns, mean±SEM) | Half-life (ns, mean±SEM) |
|---|---|---|---|---|---|
| WT | AUG | 8/8 | 5217 | 0.64 ± 0.075 | 0.23 ± 0.027 |
| WT | CUG | 8/8 | 3613 | 0.23 ± 0.026 | 0.087 ± 0.0071 |
| WT | GUG | 5/8 | 129 | 0.25 ± 0.096 | 0.11 ± 0.035 |
| WT | UUG | 8/8 | 5897 | 0.31 ± 0.056 | 0.12 ± 0.020 |
| WT | ACG | 8/8 | 2191 | 0.57 ± 0.16 | 0.24 ± 0.077 |
| WT | AUC | 8/8 | 1820 | 0.60 ± 0.054 | 0.22 ± 0.023 |
| N30K | AUG | 4/8 | 1490 | 0.21 ± 0.0091 | 0.088 ± 0.0048 |
| N30Q | AUG | 7/8 | 1064 | 0.54 ± 0.040 | 0.22 ± 0.022 |
| G29AG31A | AUG | 8/8 | 2542 | 0.43 ± 0.11 | 0.14 ± 0.025 |

**Supplementary Table 4: Mean binding lifetimes of low- and high-FRET eIF5 binding events.** Provided as a separate file (`Supplementary Table 4.xlsx`). The number of complexes and binding events analyzed, the calculated means, and 95 % C.I. are collated in this table.

**Supplementary Table 5:**
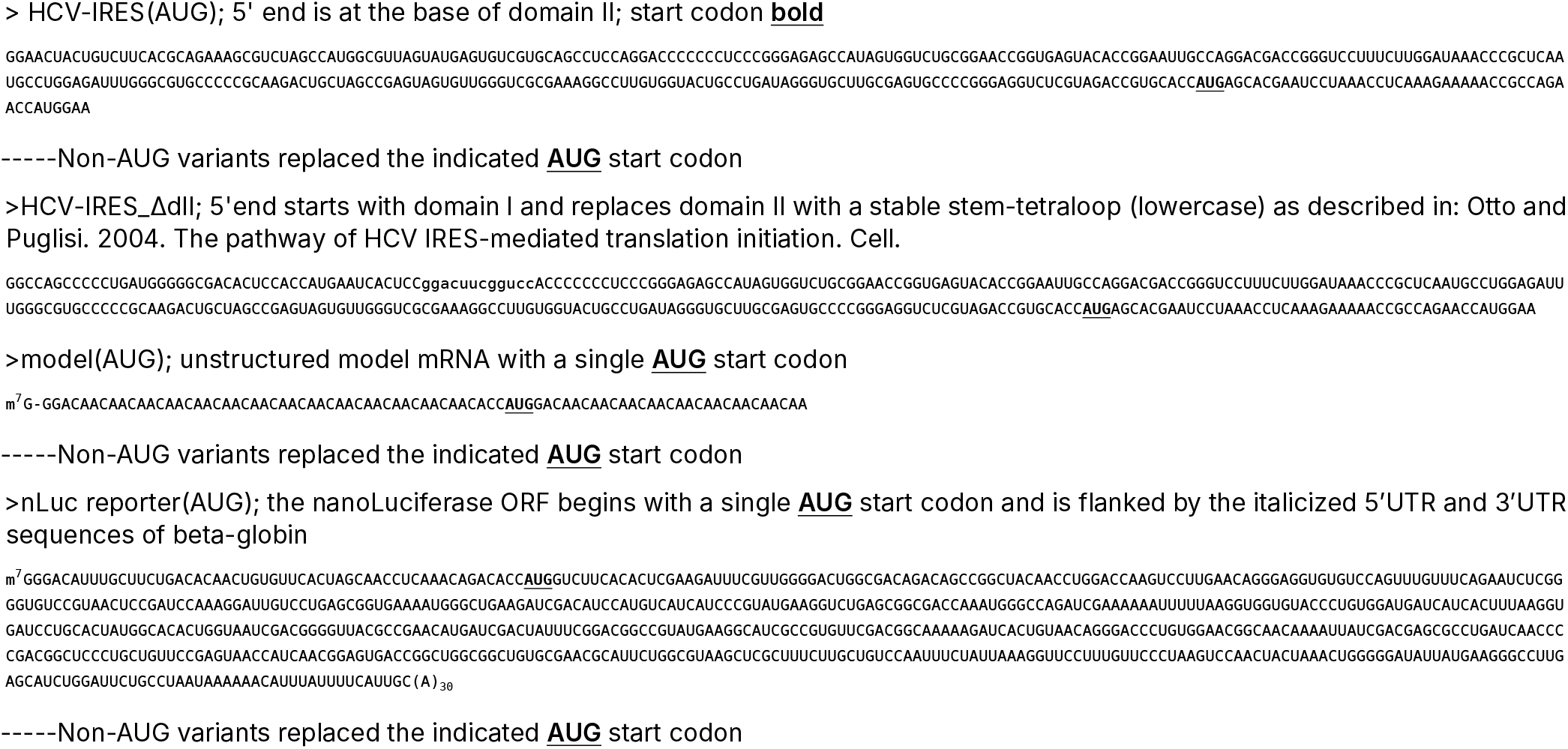
Sequences of RNAs used in this study.

**Supplementary Table 6:**
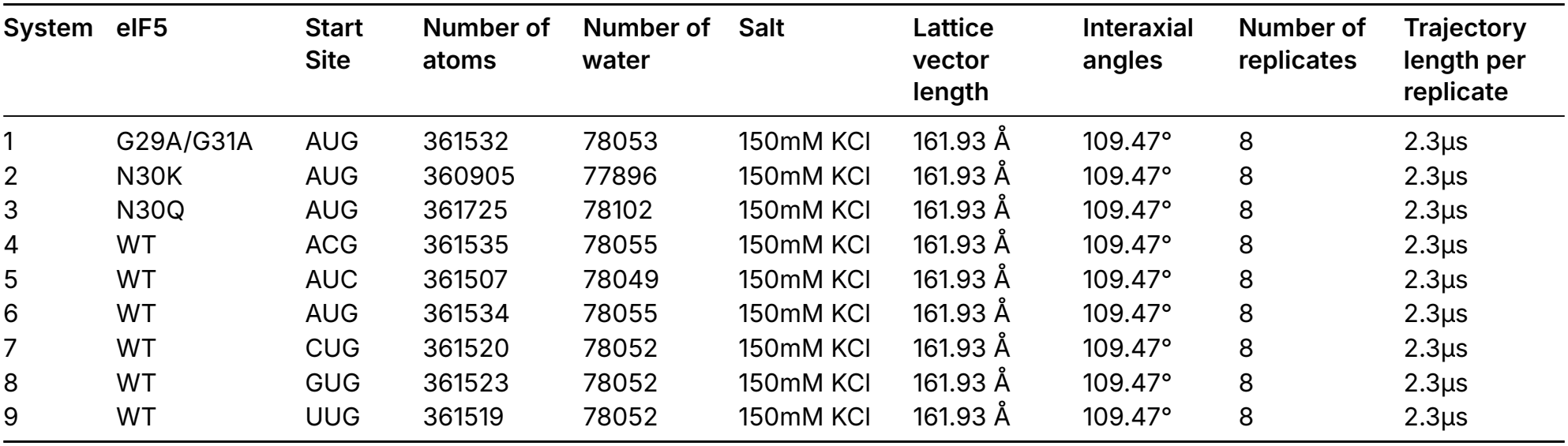
Molecular dynamics systems.

| System | eIF5 | Start Site | Number of atoms | Number of water | Salt | Lattice vector length | Interaxial angles | Number of replicates | Trajectory length per replicate |
|---|---|---|---|---|---|---|---|---|---|
| 1 | G29A/G31A | AUG | 361532 | 78053 | 150mM KCl | 161.93 Å | 109.47° | 8 | 2.3µs |
| 2 | N30K | AUG | 360905 | 77896 | 150mM KCl | 161.93 Å | 109.47° | 8 | 2.3µs |
| 3 | N30Q | AUG | 361725 | 78102 | 150mM KCl | 161.93 Å | 109.47° | 8 | 2.3µs |
| 4 | WT | ACG | 361535 | 78055 | 150mM KCl | 161.93 Å | 109.47° | 8 | 2.3µs |
| 5 | WT | AUC | 361507 | 78049 | 150mM KCl | 161.93 Å | 109.47° | 8 | 2.3µs |
| 6 | WT | AUG | 361534 | 78055 | 150mM KCl | 161.93 Å | 109.47° | 8 | 2.3µs |
| 7 | WT | CUG | 361520 | 78052 | 150mM KCl | 161.93 Å | 109.47° | 8 | 2.3µs |
| 8 | WT | GUG | 361523 | 78052 | 150mM KCl | 161.93 Å | 109.47° | 8 | 2.3µs |
| 9 | WT | UUG | 361519 | 78052 | 150mM KCl | 161.93 Å | 109.47° | 8 | 2.3µs |

